# Alphaviral capsid proteins inhibit stress granule assembly via competitive RNA binding with G3BP1

**DOI:** 10.64898/2025.12.01.691496

**Authors:** Yun Zhang, Yi Liu, Zhiying Yao, Haolong Lai, Xiaoxin Chen, Ziqiu Wang, Yiqiong Bao, Tingting Li, Xiaoming Zhou, Xiabin Chen, Peiguo Yang

## Abstract

Viral infection is one of the conditions that induces stress granule (SG) formation, a cellular defense mechanism that exerts antiviral effects. To counteract this host response, viruses have evolved a broad spectrum of strategies to inhibit SG formation. However, the molecular mechanisms underlying SG inhibition remain poorly understood. The nucleocapsid proteins play a critical role in virus replication and host interaction. Here, using Semliki Forest Virus (SFV) as a model, we uncover the function of the alphavirus nucleocapsid in SG inhibition. This inhibitory function depends on oligomerization mediated by an N-terminal α-helix and with a positively charged intrinsically disordered region (IDR). We show that SFV capsid directly competes with G3BP1 for RNA binding, thereby disrupting G3BP1-RNA liquid-liquid phase separation (LLPS) *in vitro* and SG assembly in cells. This mechanism is conserved across the alphavirus family but is not shared by the nucleocapsid of SARS-CoV-2 or other endemic viruses examined. Notably, expression of a peptide from SFV capsid is sufficient to inhibit SG formation induced by Amyotrophic Lateral Sclerosis (ALS)-associated mutations, suggesting potential therapeutic applications. Our findings reveal mechanistic insight into SG modulation by the viral capsid protein and provide a possible bioengineering tool for probing SG dynamics in health and disease.

## 1. Introduction

Stress granules (SGs) are dynamic biomolecular condensates composed of untranslated mRNA and a variety of RNA-binding proteins (RBPs)^1–3^. Untranslated messenger ribonucleoprotein (mRNP) complexes condense into SGs through a network of protein-protein, protein-RNA, and RNA-RNA interactions between individual mRNPs^4,5^. Key proteins for promoting RNA condensation and SG formation are G3BP1 and its paralog G3BP2, which dimerize through an N-terminal NTF2-like (NTF2L) domain and use C-terminal RNA-binding domains to crosslink RNAs together to form SGs^6^. SGs play critical roles in cellular stress response and have been implicated in diverse pathological conditions, including tumorigenesis, neurodegenerative diseases, and viral infections^7–10^. Over the past decade, there has been a growing interest in elucidating the mechanisms and functions of SGs. Given their role in disease pathogenesis, modulating SG formation and dynamics has emerged as a promising therapeutic strategy, underscoring the need for effective SG regulators, including small molecules, peptides, and proteins.

In response to viral infection, host cells activate stress response pathways to limit viral replication. One key defense mechanism involves translational suppression, leading to the formation of SGs^11^. G3BP1/2 restrict the replication of numerous viruses, prompting many viruses to evolve strategies to antagonize these proteins and suppress SG formation^12,13^. The viral capsid protein plays at least three essential roles in the viral life cycle: selective packaging of the viral genome, protection of the nucleic acid from degradation, and facilitation of genome delivery into the host cytoplasm^14^. For instance, the nucleocapsid (N) protein of severe acute respiratory syndrome coronavirus 2 (SARS-CoV-2) inhibits SG formation by directly interacting with G3BP1/2^15^. Similarly, our previous work and recent studies have shown that the SFV non-structural protein 3 (nsP3) disrupts SG assembly by sequestering G3BP2^11^.

The Alphavirus genus comprises more than 30 species that infect a wide range of hosts and are transmitted by arthropod vectors^16,17^. These viruses cause a spectrum of clinical manifestations in humans and animals, including rash, arthritis, joint pain, and encephalitis, contributing to a significant global health burden^18,19^. Decades of research have led to a detailed understanding of alphavirus structure, replication cycle, and host interactions^20–22^, yet the precise functions of several viral proteins remain incompletely defined. Mosquito-borne alphaviruses are traditionally classified into two groups based on disease phenotype: arthritogenic and encephalitic viruses^23^. Alternatively, they can be categorized by geographic origin, Old World and New World alphaviruses, where Old World members typically induce arthritic symptoms, whereas New World viruses are more commonly associated with encephalitis and higher mortality rates^18,20,24^. Semliki Forest virus (SFV), an Old World alphavirus^25^, causes only mild febrile illness in humans but is highly neurovirulent in rodents, making it a valuable model system for studying viral replication, virus-host interactions, and innate immune responses^26–28^. SFV is a small (∼70 nm in diameter), enveloped virus with a nucleocapsid core comprising 240 copies of capsid protein encapsulating a positive-sense, single-stranded RNA genome of ∼11.8 kb. The genome encodes four non-structural proteins (nsP1, nsP2, nsP3, and nsP4), which are translated as a single polyprotein (P1234)^16,29,30^, and five structural proteins, translated as a single polyprotein (C-E3-E2-6K-E1)^31^. SFV has evolved several mechanisms to counteract the host innate immune pathway for efficient viral replication. Previous works showed that SFV nsP3 can interact with various host proteins and block SG formation^32–35^. Whether other viral components are also involved in stress granule modulation is less clear.

Here, we identify a distinct mechanism of SG modulation mediated by the alphavirus capsid protein, which differs from the SG-inhibitory function of the SARS-CoV-2 N protein. Using SFV as a model, we demonstrate that oligomerization of a positively charged tract in the N-terminal region of the capsid protein robustly suppresses SG formation under various stress conditions. Specifically, a lysine-rich motif within the N-terminus is essential for this inhibitory activity. Adjacent to this motif lies a short hydrophobic α-helix that mediates capsid oligomerization, thereby increasing the valency of lysine residues and enhancing SG suppression. Mechanistically, the lysine-rich capsid competes with G3BP1 for mRNA binding and shifts the phase separation threshold of G3BP1-RNA complexes, thereby preventing SG assembly. An SFV mutant lacking the α-helix exhibits significantly impaired replication, underscoring the functional importance of this domain. Furthermore, SG disassembly coincides with SFV replication in infected cells, revealing an additional viral strategy to counteract the antiviral functions of SGs. Collectively, our findings uncover a novel mechanism of SG modulation by an alphaviral capsid protein and establish a new platform for engineering SG dynamics in the context of viral pathogenesis and therapeutic development.

## 2. Results

### 2.1. Alphavirus nucleocapsid inhibits stress granule formation

To systematically investigate the impact of viral proteins on host stress granule formation during alphavirus infection, we individually expressed each open reading frame (ORF) from the model alphavirus, Semliki Forest Virus (SFV). SFV encodes nine ORFs, comprising four nonstructural proteins (nsPs) and five structural proteins, including the capsid and envelope proteins (Figure 1A). We utilized sodium arsenite (SA)-induced SG formation in U2OS cells as a model system. Among the expressed proteins, nsP3 colocalized with SGs marked by G3BP1, consistent with its role in viral condensate formation (Yi Liu et al., unpublished). Notably, the capsid N protein strongly inhibited SG assembly, whereas the remaining seven SFV-encoded proteins exhibited no significant effect (Figure 1B). Quantitative analysis of SG area and number revealed that the specific effect of N in SG (Figure 1C). This inhibitory effect was not specific to SA stress stimuli. SFV N protein also robustly inhibited heat shock-induced SG formation (Figure 1D). However, it did not impair SG formation triggered by osmotic stress (Figure 1E).

**Figure 1.**
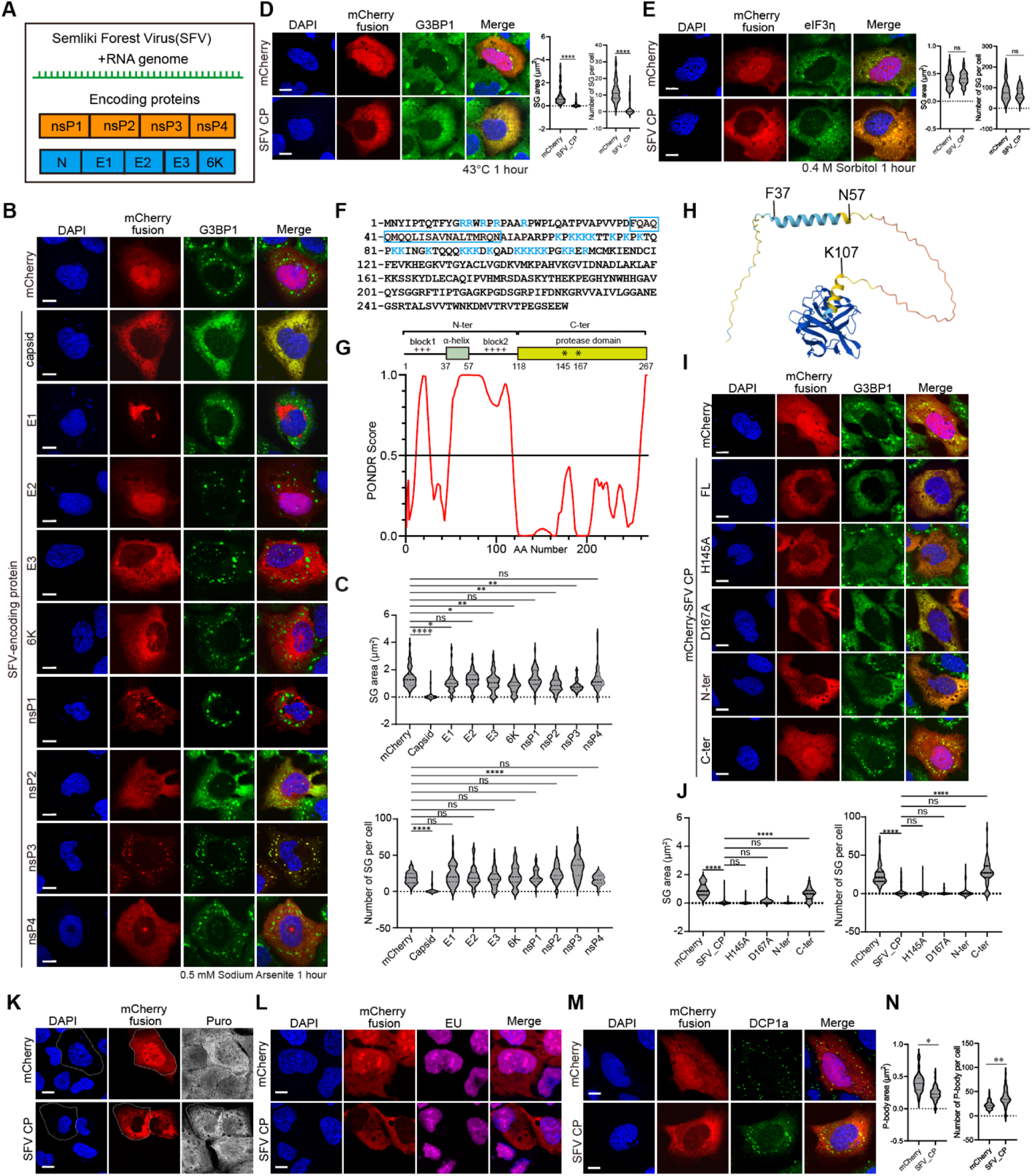
SFV capsid inhibits stress granule formation. (A) Diagram of SFV genome and encoding proteins. (B) Immunofluorescent analysis of mCherry-tagged SFV encoding proteins and endogenous G3BP1 in 0.5 mM SA-treated U2OS cells. Scale bar, 10 μm. (C) Quantification of SG area and number in (B). Data are plotted as minimum to maximum; lines indicate the first quartile (lower), median, and the third quartile (upper) with n≥50 for each sample (one-way ANOVA with Mixed-effects analysis). *p < 0.05, **p < 0.01, ****p < 0.0001, ns is not significant. Data are from one representative experiment for two independent experiments. (D) Immunofluorescence of mCherry-SFV capsid and endogenous G3BP1 in heat-stressed U2OS cells (left), with corresponding quantification of SG area and number (right). Scale bar, 10 μm. Data are plotted as minimum to maximum; lines indicate the first quartile (lower), median, and the third quartile (upper) with n≥50 for each sample (one-way ANOVA with Mixed-effects analysis). ****p < 0.0001, ns is not significant. Data are from one representative experiment for two independent experiments. (E) Immunofluorescence of mCherry-SFV capsid and endogenous eIF3η in 0.4 M sorbitol-stressed U2OS cells (left), with corresponding quantification of SG area and number (right). Scale bar, 10 μm. Data are plotted as minimum to maximum; lines indicate the first quartile (lower), median, and the third quartile (upper) with n≥50 for each sample (one-way ANOVA with Mixed-effects analysis). ns is not significant. Results of one representative experiment for two independent experiments are shown. (F) The amino acid sequence of SFV Capsid protein. The positively charged amino acids in the N-terminal of Capsid are labeled in blue, and the α-helix is marked with a box. (G) Diagram and PONDR analysis of SFV capsid protein. (H) Alphafold prediction of the structure of SFV capsid. (I) Immunofluorescence of SA-induced SG formation in U2OS cells expressing mCherry-SFV capsid truncations and protease-dead mutants. Scale bar, 10 μm. (J) Quantification of SG area and number in (I). Data are plotted as minimum to maximum; lines indicate the first quartile (lower), median, and the third quartile (upper) with n≥50 for each sample (one-way ANOVA with Mixed-effects analysis). ****p < 0.0001, ns is not significant. Results of one representative experiment for two independent experiments are shown. (K) Immunofluorescent imaging of puromycin labeling in U2OS cells expressing mCherry control and mCherry-SFV capsid. A representative mCherry or mCherry-SFV capsid cell was outlined in comparison to surrounding non-expressed cells. Scale bar, 10 μm. (L) Confocal imaging of EU-labeled nascent RNA in U2OS cells expressing mCherry and mCherry-SFV capsid. Scale bar, 10 μm. (M) Immunofluorescence of P-body marked with DCP1a in mCherry-SFV capsid expressing U2OS cells. Scale bar, 10 μm. (N) Quantification of P-body area and number in (M). Data are plotted as minimum to maximum; lines indicate the first quartile (lower), median, and the third quartile (upper) with n≥50 for each sample (one-way ANOVA with Mixed-effects analysis). *p < 0.05, **p < 0.01. Results of one representative experiment for two independent experiments are shown.

To identify the molecular determinants underlying SG inhibition, we generated a series of truncations and point mutations in the SFV N protein. The SFV capsid encodes a 267aa protein composed of an N-terminal disordered region and a C-terminal folded protease domain (Figure 1F,G). The N-ter is predicted to be disordered and enriched in positively charged residues (Figure 1G,H). To assess whether proteolytic activity contributes to SG suppression, we expressed a catalytically inactive mutant by introducing the H145A and D167A mutations into the N protein (Figure 1I,J). The protease-deficient SFV N protein can still inhibit SG formation in cells (Figure 1I,J), indicating that SG suppression is independent of protease function. By creating a series of N protein truncations and deletion mutants, we found that the N-terminal IDR alone is sufficient to inhibit SG formation (Figure 1I,J). Puromycin labeling of nascent peptides and EU-based detection of nascent RNA transcripts indicated that there is no global translational or transcriptional shutdown in the presence of N protein expression (Figure 1K,L). This suggested that global translational or transcriptional shutdown does not account for SG inhibition. Furthermore, this effect is specific to stress granules, as P-bodies formed normally in cells expressing SFV N protein (Figure 1M,N).

Collectively, these findings demonstrate that the SFV capsid protein exerts a potent and selective inhibitory effect on SG assembly.

### 2.2. N-terminal positively charged IDR mediates SG inhibition

The N-terminal region contains two positively charged segments (Figure 2A). Deletion of the second, but not the first, positively charged region abolished the inhibitory effect on SG formation, indicating that the second block is the critical determinant (Figure 2B-D). However, the second charged block alone is insufficient to inhibit SG assembly, suggesting a requirement for adjacent sequences. A predicted α-helix is located between the two charged regions (Figure 1H). Only the combination of this α-helix with the second positively charged block, and not with the first, is sufficient to inhibit SG formation. Collectively, these results identify a specific fragment of the SFV capsid, comprising the α-helix and adjacent positively charged residues, as a functional modulator of stress granule dynamics.

**Figure 2.**
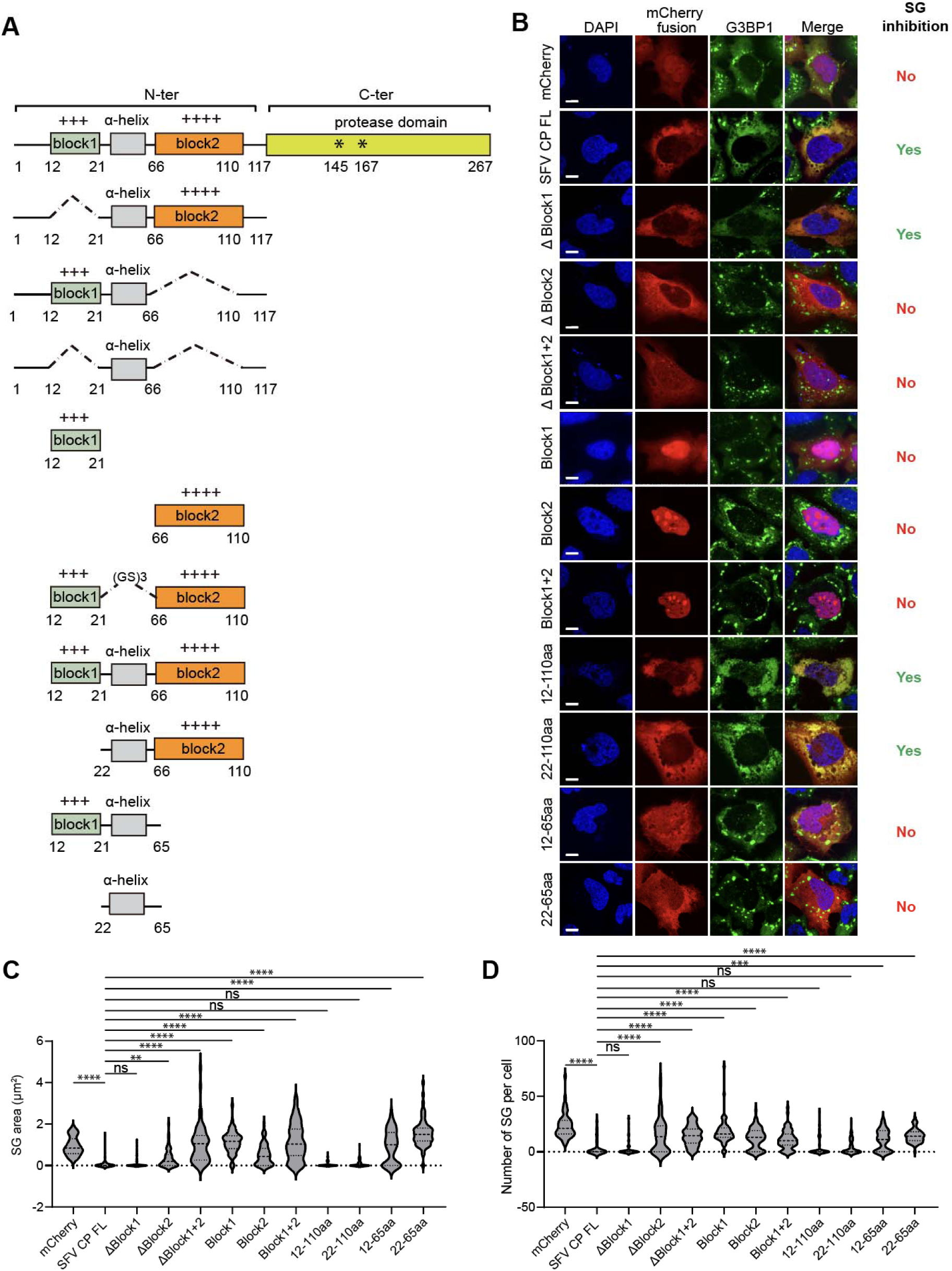
Identification of a minimal fragment of SFV capsid required for SG inhibition. (A) Schematic structures of SFV capsid N-terminal variants. (B) Immunofluorescent analysis of SA-induced SG formation in U2OS cells expressing the N-terminal variants in (A). Scale bar, 10 μm. (C, D) Quantification of SG area and number in (B). Data are plotted as minimum to maximum; lines indicate the first quartile (lower), median, and the third quartile (upper) with n≥50 for each sample (one-way ANOVA with Mixed-effects analysis). **p < 0.01, ***p < 0.001, ****p < 0.0001, ns is not significant. Data are from one representative experiment for at least two independent experiments.

### 2.3. The oligomerization motif is required for SG modulation

To characterize the feature of the helix-forming region, we performed N-terminal truncations to map the minimal motifs required for SG formation (Figure 3A,B). Starting from the above mapped region, we sequentially removed three residues from the N-terminus (Figure 3B). We observed that the fragment containing residues 40-65aa of SFV N retained the ability to inhibit SG formation in cells, whereas the 43-65aa fragment did not (Figure 3C-E). This minimal region coincides with the predicted α-helical domain (Figure 1H). To validate the helical propensity of this motif, we synthesized a peptide corresponding to residues 35-64aa of SFV capsid protein. Circular dichroism (CD) spectroscopy revealed a spectrum characteristic of α-helix formation (Figure 3F). The helix is predicted to adopt a coiled-coil (CC) conformation, a structural motif commonly associated with proteinoligomerization^36–38^. To disrupt this potential coiled-coil structure, we introduced leucine-to-proline substitutions (2LP mutant). CD analysis showed that the 2LP mutant exhibited a marked reduction in helical content (Figure 3F). AlphaFold structural predictions further supported an oligomerization-prone conformation for the wild-type α-helix, which was significantly perturbed by the 2LP mutations (Figure 3G). These findings suggest that the α-helix mediates oligomerization of this positively charged peptide, potentially facilitating efficient SG modulation. Consistent with this hypothesis, introducing 2LE or 2LP mutations into the full-length N protein significantly impaired its ability to inhibit SG formation in cells (Figure 3H-J). To directly evaluate whether oligomerization is responsible for this functional defect, we introduced cysteine residues adjacent to the helix-forming region to enable disulfide bond-mediated cross-linking^39^. The cysteine substitution did not impair the SG-inhibitory activity of the peptide (Figure S1A). Under non-reducing conditions, the cysteine-containing construct formed dimers, which were abolished upon DTT treatment (Figure 3K). No cross-linking was detected in the absence of cysteine substitution. Notably, the construct containing two cysteine residues formed higher-order oligomers, including trimers and tetramers (Figure 3K), indicating that the N-terminal IDR of SFV N can adopt an oligomeric state in cells.

**Figure 3.**
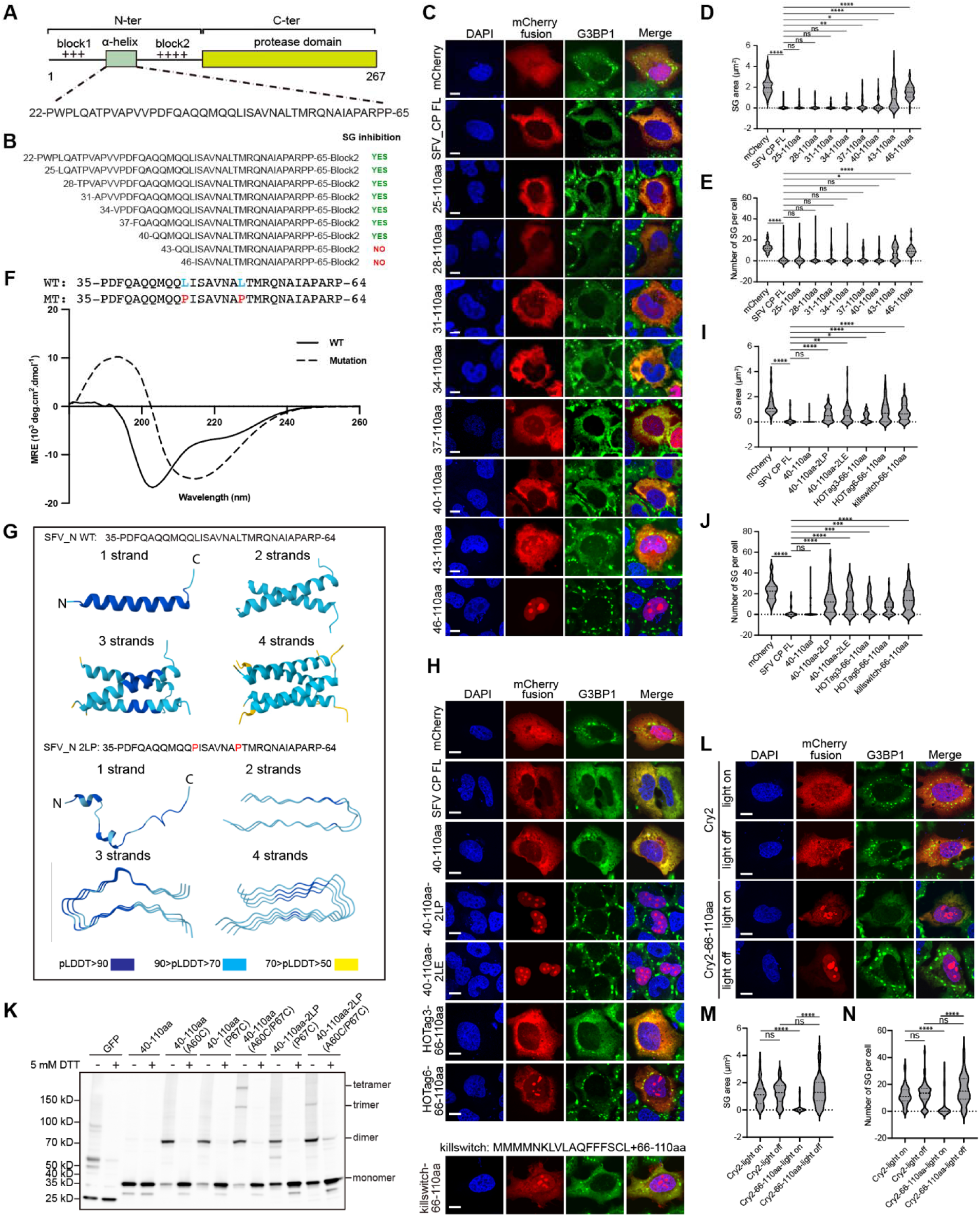
Oligomerization mediated by a helix within SFV capsid is required for SG inhibition. (A) Schematic showing the sequence of the α-helix between the two positively charged blocks in the N-terminal of SFV capsid. (B) Schematic of the SFV capsid 22-110 aa full-length and shorter variants. (C) Immunofluorescent analysis of SA-induced SG formation in U2OS cells expressing mCherry-tagged α-helix variants in (B). Scale bar, 10 μm. (D, E) Quantification of SG area and number related to (C). Data are plotted as minimum to maximum; lines indicate the first quartile (lower), median, and the third quartile (upper) with n≥50 for each sample (one-way ANOVA with Mixed-effects analysis). *p < 0.05, **p < 0.01, ****p < 0.0001, ns is not significant. Data are from one representative experiment for two independent experiments. (F) The amino acid sequences and the CD spectra of two peptides spanning the α-Helix 35-65 aa WT and the 2LP mutant. The mutation sites were highlighted in blue and red. (G) AlphaFold3 prediction of the potential SFV capsid α-helix WT conformation as monomer and multimer (Top) and the conformational prediction of its 2LP mutant (Bottom). pLDDT, predicted local distance difference test. (H) Immunofluorescent analysis of SA-induced SG formation in U2OS cells expressing the mCherry-tagged capsid 40-110aa variants, including the α-helix oligomerization-disruptive mutants of 2LP/2LE and the α-helix-displaced mutants of HOTag3/6/Killswitch. Scale bar, 10 μm. (I, J) Quantification of SG area and number related to (H). Data are plotted as minimum to maximum; lines indicate the first quartile (lower), median, and the third quartile (upper) with n≥50 for each sample (one-way ANOVA with Mixed-effects analysis). *p < 0.05, **p < 0.01, ***p < 0.001, ****p < 0.0001, ns is not significant. Data are from one representative experiment for two independent experiments. (K) HEK293T cells were transfected with EGFP-tagged SFV capsid 40–110aa or 40–110aa 2LP constructs harboring cysteine mutations, followed by Western blot analysis. (L) Immunofluorescence analysis of SA-induced SG formation in U2OS cells expressing the CRY2-helix-replaced SFV capsid 40–110aa mutant, with Cry2 alone as the negative control, in the presence or absence of blue light stimulation. Scale bar, 10 μm. (M, N) Quantification of SG area and number related to (L). Data are plotted as minimum to maximum; lines indicate the first quartile (lower), median, and the third quartile (upper) with n≥50 for each sample (one-way ANOVA with Mixed-effects analysis). ****p < 0.0001, ns is not significant. Data are from one representative experiment for two independent experiments.

To further investigate the role of oligomerization, we used a domain-swap strategy by replacing the helix-forming motif with synthetic oligomerization domains, HOTag3 (hexameric) and HOTag6 (tetrameric). Artificial oligomerization of the positively charged region partially restored SG inhibition (Figure 3H-J), with the extent of rescue correlating with the oligomeric state; HOTag3 exhibited a stronger effect than HOTag6. Furthermore, when we fused the region to Cry2, a protein known to oligomerize upon blue light exposure, we observed enhanced SG inhibition specifically upon illumination (Figure 3H-J), supporting a role for higher-order oligomerization in functional activity. As negative controls, we tested Cry2 without fusion to the SFV N peptide and without blue light stimulation. Neither condition affected SG formation (Figure 3L-N). We also examined the Killswitch tag, a recently identified self-interacting peptide capable of modulating biomolecular condensates^40^. However, it showed no significant effect on SG modulation compared to HOTag and Cry2 fusions (Figure 3H-J), likely due to its weaker oligomerization propensity.

Notably, the wild-type peptide localized predominantly to the cytoplasm (Figure 3H, third row), whereas the 2LP mutant accumulated in the nucleus and was enriched within the nucleus (Figure 3H, fourth row). This nuclear and nucleolar localization aligns with previously reported nuclear localization signals (NLS) and nucleolar targeting properties^41,42^. This pattern was recapitulated in the truncation experiment (Figure 3C), where disruption of the α-helix correlated with nuclear accumulation. Consistently, the HOTag3 and HOTag6 fusion constructs (Figure 3H, sixth and seventh row), which restored SG inhibition, also exhibited increased cytoplasmic localization.

In summary, truncation analysis, cysteine-mediated cross-linking, and domain-swap experiments demonstrate that the α-helical motif is essential for SG inhibition by promoting oligomerization. Subcellular localization further correlates with functional activity, underscoring the importance of cytoplasmic retention and multivalency in mediating this regulatory effect.

### 2.4. Capsid N protein localization during SFV replication correlates with SG inhibition

Transient transfection results in variable cellular concentrations of SFV N protein in cells, enabling estimation of the threshold concentration required for SG inhibition. By generating a standard curve using recombinant GFP-N protein *in vitro*, we converted relative GFP-N fluorescence intensity into absolute protein concentrations (Figure S1B,C). The threshold concentration of SFV N necessary to inhibit SG formation was determined to be approximately 0.41 μM (Figure S1D,E). Western blot analysis of SFV N expression at various time points post-infection, combined with a standard curve for SFV N, revealed that intracellular N protein levels reach approximately 2.4 μM by 8 hours post-infection (hpi) (Figure S1F), suggesting that the SFV N accumulated early during infection to concentrations sufficient to inhibit SG assembly.

To examine capsid localization during virus infection, we generated a recombinant SFV virus with a GFP11 tag inserted near the N-terminus of the capsid protein (Figure 4A), a site previously reported to be permissive for viral infectivity^43^. To minimize potential perturbations caused by the tag, we generated an antibody using bacteria-expressed capsid proteins. The antibody effectively recognizes the recombinant capsid protein expressed in cells (Figure 4B). Using the GFP11-tagged SFV, we observed that the capsid protein is predominantly diffusely distributed in the cytoplasm. However, in approximately 20–50% of infected cells, punctate foci were detected (Figure 4C). We further examined the relationship between N foci and the SFV replication complex marked by nsP3 in cells and found that more cells exhibited nsP3 foci than those with capsid foci (Figure 4D). In cells with both capsid and nsP3 foci, these structures frequently colocalized (Figure 4D). Previous studies, including our own, have demonstrated that the alphaviral nsP3 promotes virus replication by forming viral condensates and localizes to membrane-bound spherules^44,45^. Our recombinant SFV system enables us to track the dynamics of the capsid during infection and suggests a tight relationship between capsid foci and the viral replication compartment.

**Figure 4.**
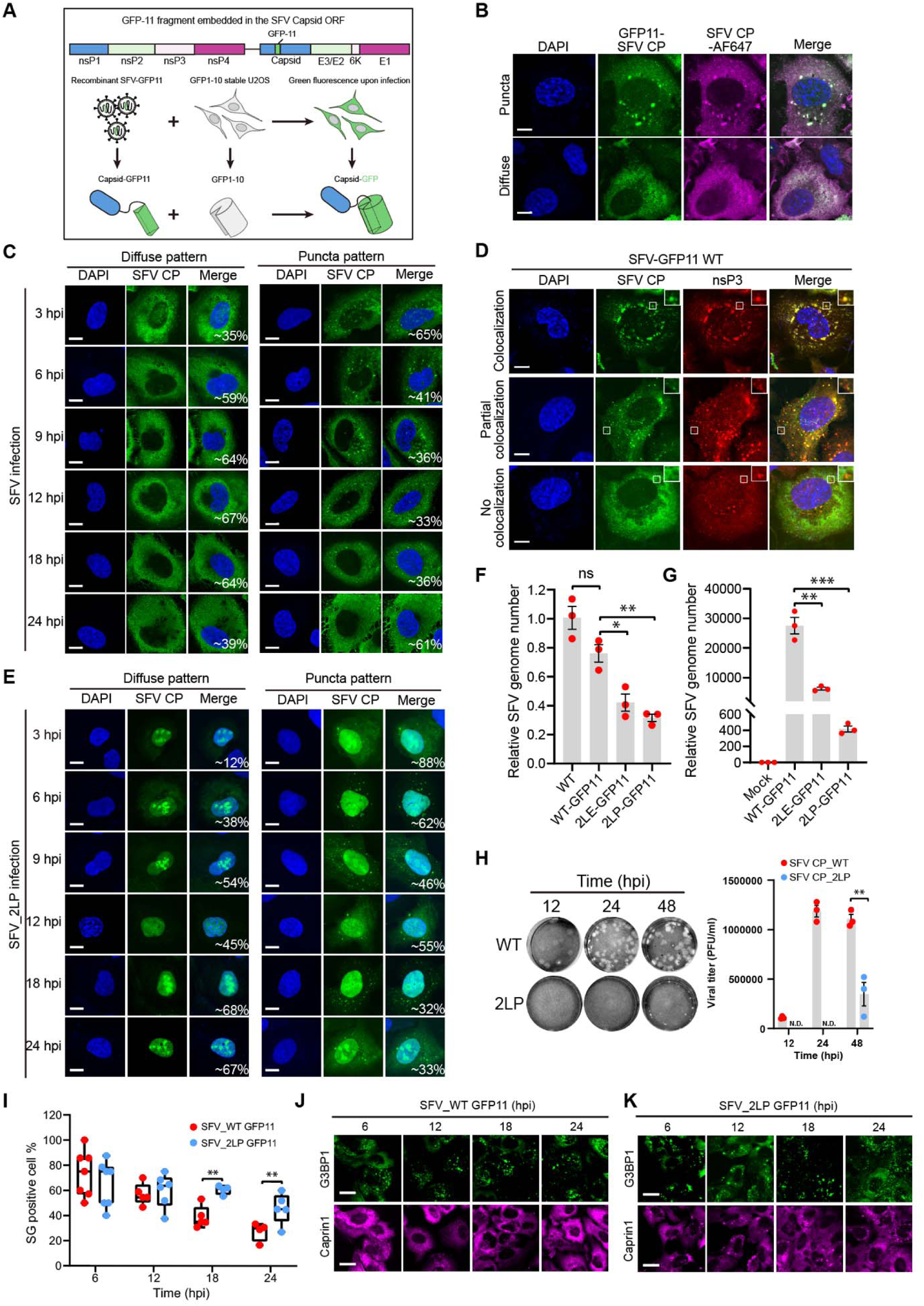
Disruption of α-Helix of capsid dampens SFV replication and SG inhibition activity in cells. (A) Schematic of visualizing SFV capsid during viral infection with a recombinant SFV containing the GFP11 tag in-frame with the capsid ORF. Infection of the GFP1-10 stable U2OS cells with the GFP11-SFV enabled the reassembly of a functional GFP and a fluorescent readout. The diagram was created with BioRender.com. (B) Confocal imaging of SFV capsid in GFP-11 SFV infected U2OS cells co-stained with SFV capsid antibody. Representative patterns of diffuse and puncta were shown. Scale bar, 10 μm. (C) Confocal imaging of the capsid expression pattern in WT SFV-infected U2OS cells at the indicated time points. Percentage of the patterns was quantified by examining ≥50 cells for each time point from one of the two independent experiments. Scale bar, 10 μm. (D) Confocal imaging of SFV capsid in GFP-11 SFV infected U2OS cells co-stained with a SFV nsP3 antibody. Three representative patterns are detected: full co-localization of capsid with nsP3 foci (top); partial co-localization of capsid with nsP3 foci (middle); diffuse capsid without substantial co-localization with nsP3 foci (bottom). Scale bar, 10 μm. (E) Confocal imaging of the capsid expression pattern in α-helix 2LP-mutated SFV-infected U2OS cells at the indicated time points. Percentage of the patterns was quantified by examining ≥50 cells for each time point from one of the two independent experiments. Scale bar, 10 μm. (F) RT-qPCR analysis of SFV WT and variants replication in HEK293T cells transfected with the indicated SFV genome-encoding constructs. Data are mean ± SEM. Results of one representative experiment for three independent experiments are shown. (G) RT-qPCR analysis of SFV WT and variants replication in HEK293T cells infected with the indicated recombinant SFV at an MOI of 0.1. Data are mean ± SEM. Results of one representative experiment for three independent experiments are shown. (H) U2OS cells were infected with SFV WT and 2LP at an MOI of 0.1. Virus-containing supernatants at the indicated time points were collected to infect confluent BHK21 cells for 5 days. Left: the representative images of plaques formed by the SFV WT and 2LP at each time point; right: quantification of the growth kinetics of SFV WT and 2LP by the plaque assay. Results of one representative experiment for two independent experiments are shown. (I-K) Representative images and quantification of SG dissolution kinetics in SFV WT- and 2LP-infected U2OS cells at the indicated time points. SGs were defined as G3BP1/Caprin1 double-positive foci. For quantification, five fields of view (FOVs) with an average of 20 cells per FOV were analyzed. Representative results of one experiment for two independent experiments are shown. Data are plotted as minimum to maximum (I); lines indicate the first quartile (lower), median, and the third quartile (upper). Scale bar, 20 μm.

SGs can be observed in the early stages of SFV infection, and the ratio of SG-positive cells decreases over time (Yi Liu et al, unpublished). Given the established role of nsP3 in SG modulation, we sought to determine whether the capsid N protein also contributes to the kinetics of SG disassembly. To this end, we introduced the 2LP mutation into the GFP11-SFV construct. We observed predominant nuclear localization of the capsid during SFV-2LP infection, compared with cytoplasmic localization during WT SFV infection (Figure 4E). Despite significant impairment in replication, assessed by RT-PCR following plasmid transfection (Figure 4F) and serial virus passage (Figure 4G,H), we were able to recover viable SFV-2LP virus. Quantification of SG formation over a 24-hour infection window revealed a strong inverse correlation between diffuse cytoplasmic N protein expression and SG presence (Figure 4I,J). SFV-2LP-infected cells exhibited prolonged SG persistence, consistent with attenuated viral replication (Figure 4I,K). Thus, the impaired fitness of the SFV-2LP mutant may be partially attributed to defective SG suppression.

In summary, the SFV capsid N protein is predominantly expressed in a diffuse cytoplasmic pattern throughout the course of infection. The inverse correlation between N protein expression and SG formation supports a model in which capsid-mediated SG inhibition acts in parallel to nsP3-dependent mechanisms to counteract host antiviral responses and facilitate viral replication.

### 2.5. The valency and density of the positive charges within capsid IDR determine the SG modulation effect

To further characterize the roles of the positively charged region within SFV N in SG modulation, we generated three scrambled mutants within the positive charge region (Figure 5A). All three mutants showed robust SG inhibition, comparable to WT proteins, in both full-length and N-terminal constructs (Figure 5B,C). To test the influence of lysine patterning, we created mutants with evenly distributed lysines or clustered lysine arrangements (Figure 5D). Both the uniformly distributed and clustered mutants effectively inhibited SG formation (Figure 5E,F), indicating that the spatial arrangement of lysine residues is not critical for this function.

**Figure 5.**
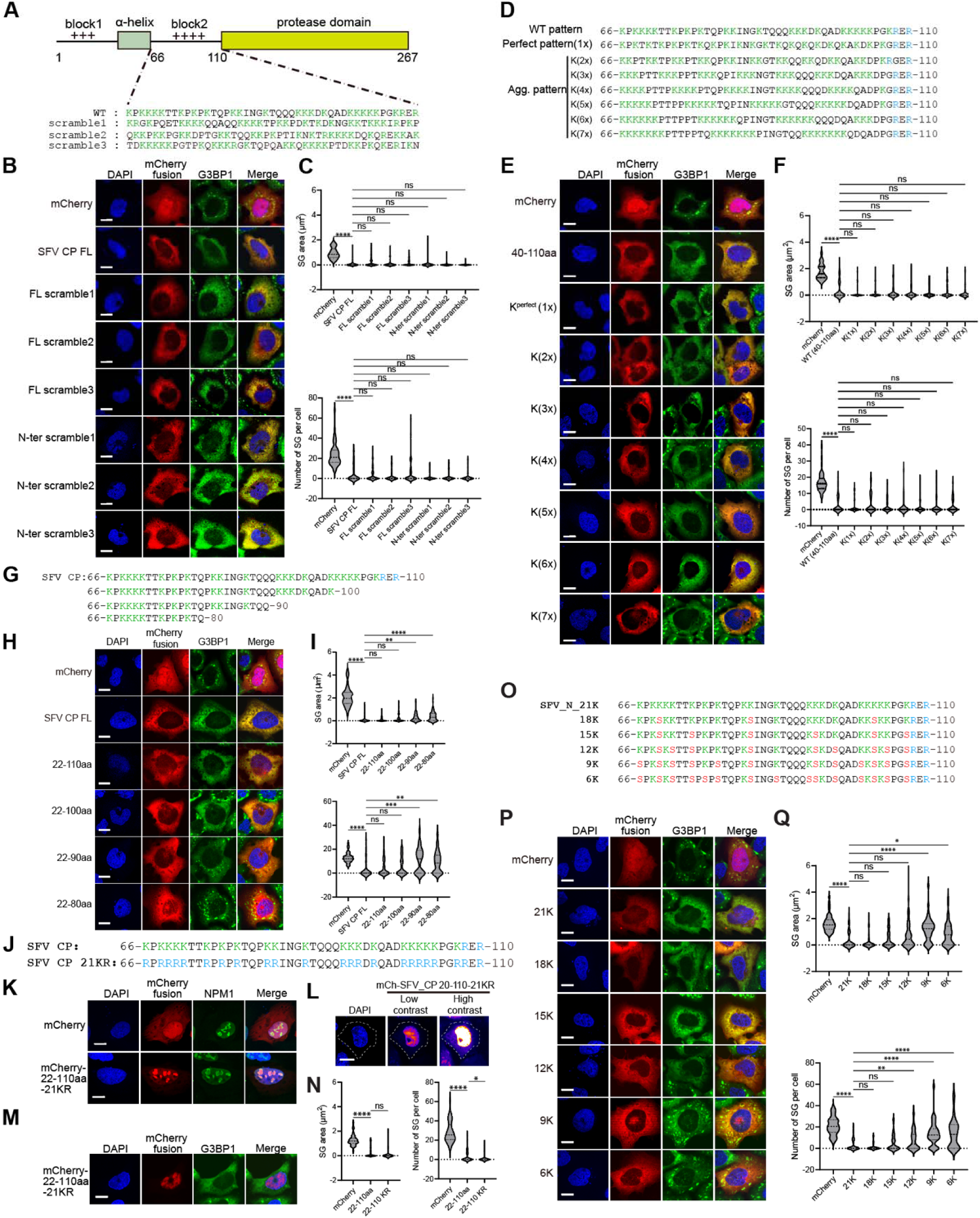
The valency and density of the positive charges are required for SG modulation. (A) Schematic of SFV capsid with positively charged block2 and 3 scramble mutants. Lysine and arginine residues in each sequence were highlighted in green. (B) Immunofluorescent analysis of SA-induced SG formation in U2OS cells expressing the mCherry-tagged block 2 scrambles in the context of full-length and N-terminal truncation. Scale bar, 10 μm. (C) Quantification of SG area and number in (B). Data are plotted as minimum to maximum; lines indicate the first quartile (lower), median, and the third quartile (upper) with n≥50 for each sample (one-way ANOVA with Mixed-effects analysis). ****p < 0.0001, ns is not significant. Data are from one representative experiment for at least two independent experiments. (D) Schematic of lysine pattern rearrangement of SFV capsid block 2. (E) Immunofluorescent analysis of SA-induced SG formation in U2OS cells expressing the indicated lysine-rearranged 40-110 aa constructs. Scale bar, 10 μm. (F) Quantification of SG area and number related to (E). Data are plotted as minimum to maximum; lines indicate the first quartile (lower), median, and the third quartile (upper) with n≥50 for each sample (one-way ANOVA with Mixed-effects analysis). ****p < 0.0001, ns is not significant. Data are from one representative experiment for at least two independent experiments. (G) Schematic of Lys/Arg valency variant of SFV capsid block 2. (H) Immunofluorescent analysis of SA-induced SG formation in U2OS cells expressing the mCherry-SFV capsid 22-110 aa constructs harboring either the WT or the lysine valency-shortened block 2 as in (G). Scale bar, 10 μm. (I) Quantification of SG area and number related to (H). Data are plotted as minimum to maximum; lines indicate the first quartile (lower), median, and the third quartile (upper) with n≥50 for each sample (one-way ANOVA with Mixed-effects analysis). **p < 0.01, ***p < 0.001, ****p < 0.0001, ns is not significant. Data are from one representative experiment for at least two independent experiments. (J) Schematic of arginine substitution (KR) of block 2 variant. (K) Confocal imaging of mCherry and mCherry-fused SFV capsid 22-110aa-KR co-stained with NPM1 antibody. (L) Representative high-contrast image of mCherry-fused SFV capsid 22-110aa-KR, showing cytoplasmic expression. Scale bar, 10 μm. (M) Immunofluorescent analysis of SA-induced SG formation in U2OS cells expressing the mCherry-SFV capsid 22-110 aa 21KR. (N) Quantification of SG area and number in (M). Data are plotted as minimum to maximum; lines indicate the first quartile (lower), median, and the third quartile (upper) with n≥50 for each sample (one-way ANOVA with Mixed-effects analysis). *p < 0.05, ****p < 0.0001, ns is not significant. Data are from one representative experiment for two independent experiments. Scale bar, 10 μm. (O) Schematic of block 2 Lys valency variants in which the positively-charged lysine was displaced by the non-charged serine to keep the total residue counts identical. (P) Immunofluorescent analysis of SA-induced SG formation in U2OS cells expressing the mCherry-SFV capsid 40-110 aa constructs harboring the indicated lysine valency as in (O). Scale bar, 10 μm. (Q) Quantification of SG area and number related to (P). Data are plotted as minimum to maximum; lines indicate the first quartile (lower), median, and the third quartile (upper) with n≥50 for each sample (one-way ANOVA with Mixed-effects analysis). *p < 0.05, **p < 0.01, ****p < 0.0001, ns is not significant. Data are from one representative experiment for at least two independent experiments.

To examine the role of lysine valency, we performed C-terminal truncations to generate a series of lysine-rich fragments of varying lengths (Figure 5G). A fragment containing 16 lysine residues exhibited SG inhibitory activity similar to the WT fragment, which contains 21 lysines and 2 arginines (Figure 5H,I). Further truncation to 11 lysines resulted in reduced inhibitory capacity, while a fragment with only 8 lysines completely lost this ability. These findings indicate a valency-dependent effect, with 11-16 lysine residues required for effective SG inhibition. The SFV N protein contains a lysine-enriched region with 21 lysines and 2 arginines within a 45-amino acid segment. To evaluate the functional interchangeability of lysine and arginine, we generated a mutant in which all lysines were substituted with arginines (Figure 5J). This arginine-rich variant displayed predominantly nucleolar localization, as confirmed by NPM1 staining (Figure 5K). Despite relatively high nuclear expression, the 21KR mutant retained a robust SG-inhibition effect (Figure 5M,N).

To maintain the overall length of the 45aa peptide while varying lysine content, we generated a series of constructs with progressively fewer lysine residues. The construct containing 12 lysines showed reduced ability in SG inhibition (Figure 5O), whereas the 15 lysines construct performed comparably to WT (Figure 5P,Q). In contrast, the 9-lysine construct completely lost SG inhibition ability. Further fine-mapping revealed that a construct with 11 lysines exhibited significantly reduced activity (Figure 5P,Q), consistent with the result from the C-terminal truncation (Figure 5G-I). Threshold concentration measurement demonstrated that the 11-lysine construct required higher expression levels for SG inhibition, with a functional transition occurring between 11 and 12 lysine residues (Figure S1G). Although subtle effects of lysine distribution within the IDR cannot be fully excluded, our systematic analysis using truncation and substitution mutants indicates that the functional threshold lies between 11 and 12 lysine residues. Collectively, these results demonstrate that the valency and density of positively charged residues, rather than their specific patterning, are the primary determinants of SG inhibition.

### 2.6. SFV capsid binds host RNA and interferes with G3BP1-RNA phase separation

The primary function of alphavirus capsid N protein is to bind viral genome RNA and facilitate viral particle assembly. We investigated whether the SFV N protein can also interact with host RNAs when expressed individually or during infection. Given that mRNA release from translating ribosomes and phase separation with RBPs are the underlying mechanisms of SG formation, we hypothesized that competitive RNA binding by SFV N proteins could contribute to SG inhibition in cells. To determine whether this inhibitory effect is mediated through scaffold proteins G3BP1 and G3BP2, we performed TurboID-based proximity labeling to map the G3BP1 interactome in the presence of SFV N. No significant alterations in the G3BP1 interactome were observed upon SFV N expression (Figure S2A,C), nor following sodium arsenite (SA) treatment (Figure S2B,D).

Our previous study showed that phase separation between G3BP1 and free mRNA drives SG formation^6^. G3BP1 undergoes liquid-liquid phase separation (LLPS) with single-stranded RNA in a sequence-independent manner^6^. Using SFV nsP3 RNA as a model, we recapitulated G3BP1-RNA LLPS *in vitro* (Figure 6A,B), consistent with previous observations^46,47^. Notably, SFV N efficiently disrupted G3BP1-RNA condensate formation, indicating its ability to compete with G3BP1 for RNA binding (Figure 6C,D). A similar inhibitory effect was observed in the G3BP1–poly(A) RNA system (Figure 6E).

**Figure 6.**
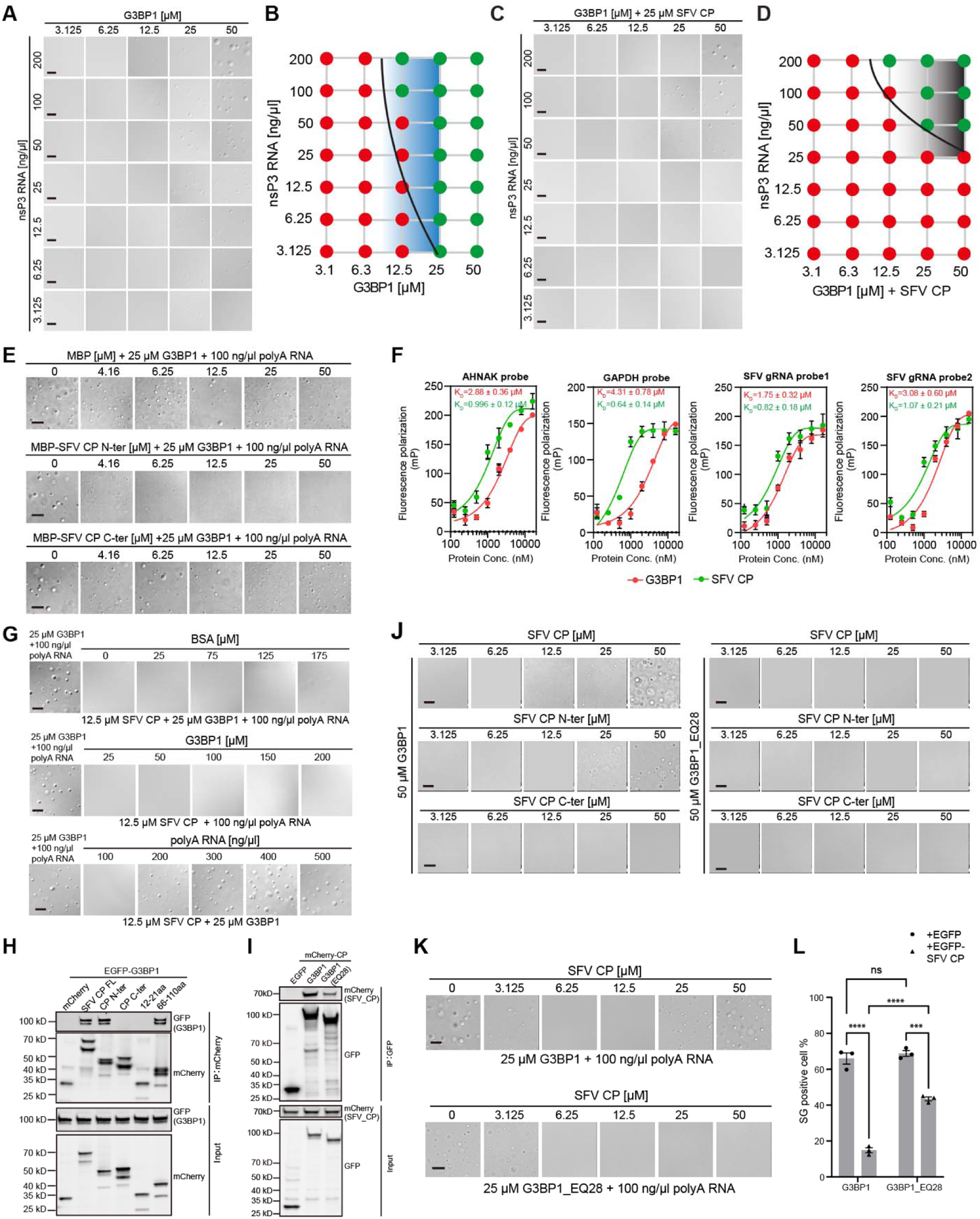
SFV Capsid inhibits G3BP1-RNA phase separation via RNA competition. (A, B) LLPS of purified G3BP1 with *in vitro*-transcribed SFV nsP3 RNA at the indicated concentrations. (B) The phase diagram of (A). (C) LLPS of purified G3BP1 with *in vitro*-transcribed SFV nsP3 RNA at the indicated concentrations in the presence of 25 μM SFV capsid. Scale bar, 10 μm. (D) The phase diagram of (C). (E) LLPS of 25 μM G3BP1 and 100 ng/μl polyA RNA in the presence of increasing concentrations of MBP, MBP-fused N-terminal, and C-terminal SFV capsid. Scale bar, 10 μm. (F) Fluorescence polarization (FP) assay to measure the binding of purified G3BP1 or SFV capsid to FAM-labeled host (n=2) and viral (n=2) RNA probes. Data are mean ± SEM with n=3 technical replicates in one representative experiment. (G) Excess RNA reverses capsid inhibition of G3BP1-RNA LLPS. Increasing concentrations of polyA RNA, G3BP1, and BSA were individually added into a specified capsid-G3BP1-RNA system, and the reoccurrence of LLPS was captured via DIC imaging. Scale bar, 10 μm. (H) IP-WB to map the SFV capsid domains required for G3BP1 interaction. Note: the double band for mCherry fusion is likely due to the proteolysis of mCherry during the boiling of samples for WB^84^. (I) IP-WB to test the interaction between SFV capsid and G3BP1_WT or G3BP1_EQ28. Note: the MW of G3BP1_EQ28 is consistently lower than that of G3BP1_WT on SGS-PAGE, likely due to charge neutralization, as both constructs contain the same GFP and linker sequence. (J) LLPS of 50 μM G3BP1_WT (left) or G3BP1_EQ28 (right) with full-length, N-terminal, or C-terminal SFV N protein at the indicated concentrations. Scale bar, 10 μm. (K) LLPS of 25 μM G3BP1_WT (top) or G3BP1_EQ28 (bottom) with 100ng/μL RNA in the presence of SFV capsid at the indicated concentrations. Scale bar, 10 μm. (L) G3BP1/2 dKO U2OS cells were co-transfected with EGFP or EGFP-SFV capsid, and mCherry-G3BP1 or mCherry-G3BP1_EQ28. Cells with SG were quantified in the indicated EGFP fusion-positive cells. Data are mean ± SEM with n=3 independent biological replicates. ***p < 0.001, ****p < 0.0001, ns is not significant.

Viral nucleocapsid, particularly the N-terminal region, has been reported to bind SFV viral RNA^48,49^. Given that SFV N suppresses SG assembly even in the absence of other viral components, we hypothesized that it might bind non-viral RNAs as well. To assess its capacity for non-viral RNA binding, we performed fluorescence polarization (FP) assays using four probes: two host mRNA probes and two viral RNA probes. SFV N exhibited higher binding affinity for all four probes (Figure 6F), supporting the hypothesis that SFV N competes with G3BP1 for RNA, thereby inhibiting SG nucleation. To further elucidate this mechanism of SG inhibition, we conducted *in vitro* LLPS rescue experiments by reintroducing either G3BP1 or RNA into SFV N-containing reactions, with BSA as a control. Droplet formation was restored upon addition of poly(A) RNA, whereas neither G3BP1 nor BSA rescued phase separation (Figure 6G), demonstrating that RNA competition alone is sufficient to overcome capsid-mediated inhibition of LLPS.

To investigate whether SFV N directly interacts with G3BP1, co-immunoprecipitation followed by western blotting (IP-WB) was performed in cells. Both full-length SFV N and its positively charged IDR in the N-terminus are associated with G3BP1 (Figure 6H). Given that G3BP1 contains an acidic IDR1, we tested the role of electrostatic interactions by employing a G3BP1 mutant with charge-neutralizing E28Q substitutions (G3BP1_EQ28). The interaction between SFV N and G3BP1 was significantly diminished in the EQ28 mutant (Figure 6I), indicating that electrostatic forces between the acidic IDR1 of G3BP1 and the positively charged region of SFV N contribute to their association. Furthermore, recombinant G3BP1 underwent phase separation with SFV N in a dose-dependent manner (Figure 6J, left panel). The N-terminal domain of SFV N, but not the C-terminal domain, was sufficient to drive phase separation with G3BP1 *in vitro*. Consistent with the IP results, phase separation between SFV N and the G3BP1_EQ28 mutant was impaired (Figure 6J, right panel), underscoring the importance of electrostatic interactions in this process. To model SG formation in a reconstituted system, SFV N was titrated into G3BP1–RNA LLPS reactions. Increasing concentrations of SFV N progressively suppressed droplet formation (Figure 6K, upper panel), while higher levels of SFV N led to droplet reappearance due to direct phase separation between G3BP1 and SFV N. Using the same system, SFV N effectively inhibited LLPS mediated by the G3BP1_EQ28 mutant in a dose-dependent manner (Figure 6K, lower panel), indicating that RNA competition with G3BP1 occurs independently of the acidic IDR1. However, phase separation between G3BP1 and SFV N was abolished in the EQ28 mutant, consistent with the two-component LLPS data (Figure 6J). Finally, we evaluated the ability of SFV N to inhibit SG formation driven by the G3BP1_EQ28 mutant in G3BP1/2 double knockout cells. Robust SGs formed in the presence of the EQ28 mutant. Although SFV N effectively suppressed SG assembly induced by both wild-type G3BP1 and the EQ28 mutant, the inhibitory effect was less pronounced with the mutant, as reflected by the higher proportion of cells retaining SGs (Figure 6L). The above results demonstrate a direct interaction between G3BP1 and SFV N, and reveal that SFV N can effectively compete with G3BP1 for RNA binding, inhibiting SG formation *in vitro* and in cells.

To comprehensively evaluate SFV N-host RNA interactions, we performed RNA immunoprecipitation followed by sequencing (RIP-seq) using GFP nanobodies in HEK293T cells expressing GFP-tagged SFV capsid, under basal conditions or upon SA treatment or SFV infection. Peak analysis showed differential enrichment of transcripts bound by capsid in SA-treated cells and SFV-infected cells, with 89 transcripts common to both conditions (Figure S3A,B). Within target transcripts, SFV N preferentially bound coding sequence (CDS) regions (Figure S3C). Comparison with a previously published SG-enriched transcriptome^50^ revealed significant overlap between SG-associated RNAs and those bound by SFV N under stress, suggesting that the capsid competes for a subset of G3BP1-bound mRNAs (Figure S3D,E). In contrast, minimal overlap was observed between the SFV N-bound transcriptome and that of the SARS-CoV-2 N protein^51^, indicating distinct RNA-binding specificities (Figure S3F).

Collectively, these findings demonstrate that SFV N directly competes with G3BP1 for RNA binding, disrupts G3BP1-RNA phase separation, and thereby suppresses stress granule assembly through a competitive RNA binding mechanism.

### 2.7. SG modulation is a conserved feature among alphavirus capsids

Amino acid sequence alignment of all 32 known alphavirus capsid proteins reveals a highly conserved C-terminal domain (CTD) and a more variable N-terminal domain (NTD) (Figure S4A). Despite limited overall sequence similarity, the distribution and density of positively charged residues are remarkably conserved across species. Secondary structure predictions indicate conservation of an α-helical region, even in the absence of strong sequence homology. Notably, the two critical leucine residues within the α-helix of SFV N are 100% conserved, underscoring their essential functional role during viral evolution (Figure S4B). The alignment further identifies a conserved modular architecture in alphavirus capsids, shared between New World and Old World alphaviruses, albeit with distinct amino acid compositions. A phylogenetic analysis based on capsid sequences effectively segregates the alphaviruses into New World and Old World clades (Figure S4C).

To broaden the scope of viral capsids capable of SG inhibition, we expressed three capsid proteins from New World alphaviruses, Venezuelan equine encephalitis virus (VEEV), Eastern equine encephalitis virus (EEEV), and Western equine encephalitis virus (WEEV), alongside four Old World alphavirus capsids: Ross River virus (RRV), Sindbis virus (SINV), Chikungunya virus (CHIKV), and Onyong-nyong virus (ONNV). Expression of all seven capsid proteins, representing both evolutionary groups, robustly suppressed SG formation (Figure 7A,B). Truncation analyses demonstrated that the N-terminal IDRs enriched in positively charged residues are sufficient for SG inhibition (Figure 7C,D), highlighting a conserved mechanistic role for positive charge in this process.

**Figure 7.**
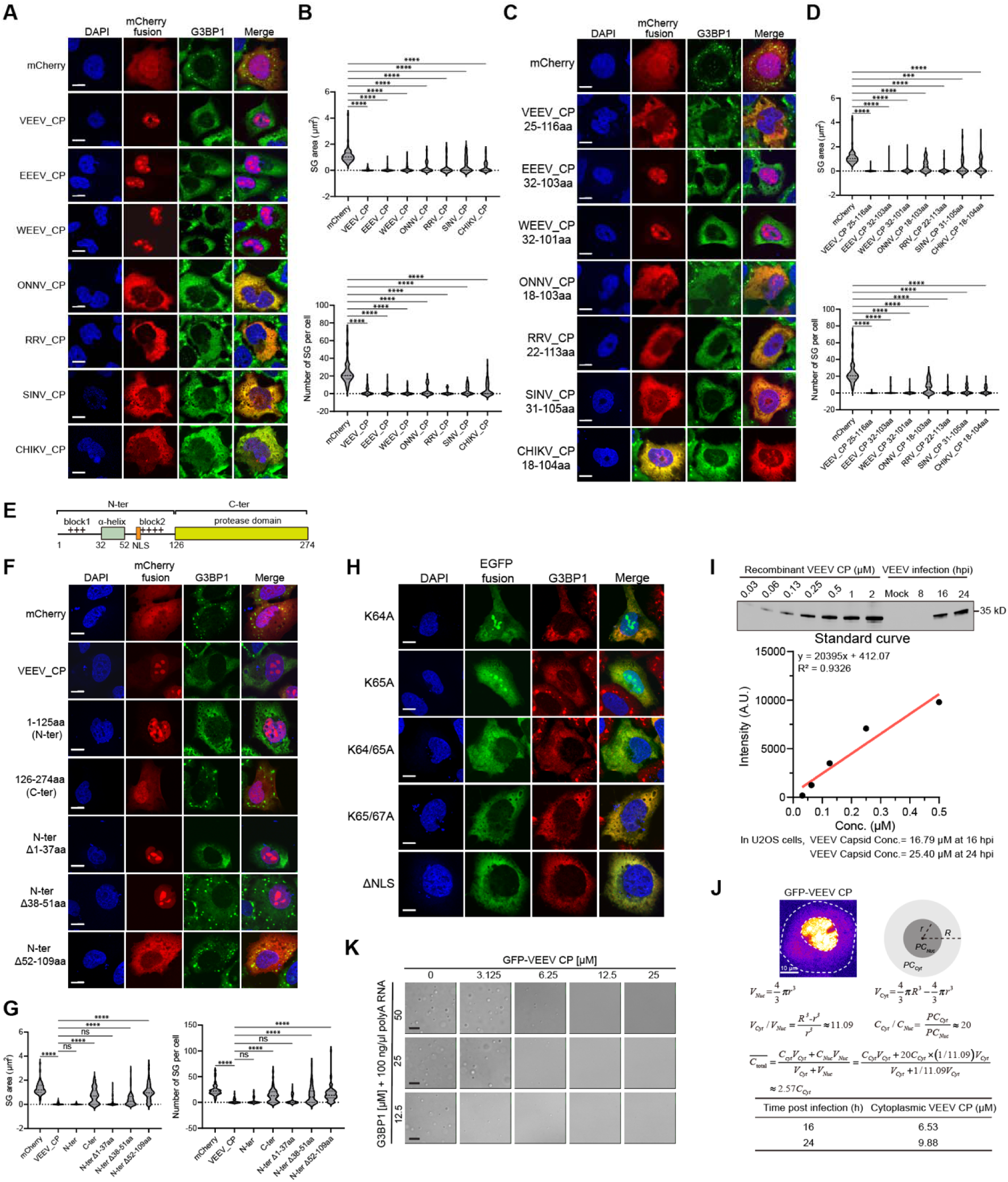
A shared feature of alphavirus capsids on SG inhibition. (A) Immunofluorescent analysis of SA-induced SG formation in U2OS cells expressing the indicated mCherry-tagged alphaviral capsids. Scale bar, 10 μm. (B) Quantification of SG area and number in (A). Data are plotted as minimum to maximum; lines indicate the first quartile (lower), median, and the third quartile (upper) with n≥50 for each sample (one-way ANOVA with Mixed-effects analysis). ****p < 0.0001. Data are from one representative experiment for two independent experiments. (C) Immunofluorescent analysis of SA-induced SG formation in U2OS cells expressing the mCherry-fused α-helix-block 2 region of the indicated alphaviral capsids. Scale bar, 10 μm. (D) Quantification of SG area and number in (C). Data are plotted as minimum to maximum; lines indicate the first quartile (lower), median, and the third quartile (upper) with n≥50 for each sample (one-way ANOVA with Mixed-effects analysis). ****p < 0.0001. Data are from one representative experiment for at least two independent experiments. (E) Schematic architecture of VEEV capsid protein. The block 2 contained a nuclear localization signal (NLS). (F) Immunofluorescent analysis of SA-induced SG formation in U2OS cells expressing the indicated mCherry-fused truncations of VEEV capsid. Scale bar, 10 μm. (G) Quantification of SG area and number in (F). Data are plotted as minimum to maximum; lines indicate the first quartile (lower), median, and the third quartile (upper) with n≥50 for each sample (one-way ANOVA with Mixed-effects analysis). ****p < 0.0001, ns is not significant. Data are from one representative experiment for at least two independent experiments. (H) Immunofluorescent analysis of SA-induced SG formation in U2OS cells expressing the indicated EGFP-tagged NLS mutants of VEEV capsid. Scale bar, 10 μm. (I) Intracellular concentration estimation of total VEEV capsid protein at 16 and 24 hpi in VEEV-infected U2OS cells (MOI=0.1). See the method section for the detailed procedure. (J) Intracellular concentration estimation of total and cytoplasmic VEEV capsid protein based on the expression ratio of nuclear over cytoplasmic GFP-VEEV N protein in U2OS cells. (K) LLPS of G3BP1 and 100 ng/μl polyA RNA in the presence of the indicated concentrations of VEEV capsid. Scale bar, 10 μm.

Strikingly, the subcellular localization of these capsid proteins diverges between the two alphavirus groups: New World capsids, exemplified by VEEV, predominantly localize to the nucleus, with strong enrichment in the nucleolus, whereas all tested Old World capsid proteins are primarily cytoplasmic (Figure 7A). Using VEEV capsid as a representative of the New World lineage (Figure 7E), we found that both the helix-forming segment and the positively charged residues are essential for SG inhibition (Figure 7F,G). Mutation of a single lysine residue partially relocalized VEEV capsid to the cytoplasm, while double mutations or deletion of a predicted nuclear localization signal (NLS) within the positively charged region were sufficient to shift the majority of capsid expression to the cytoplasm (Figure 7H). In all cells expressing cytoplasm-localized VEEV capsid variants, SG formation remained strongly suppressed. These findings suggest an evolutionarily divergent strategy in subcellular targeting between New World and Old World alphavirus capsids. We further synthesized the N-terminal domains from all remaining 24 alphavirus capsids, including the helix-forming region and adjacent positively charged segments (Figure S4A), and confirmed that each is capable of inhibiting SG assembly (Figure S5A), reinforcing the functional conservation of this mechanism. Given the high degree of conservation among alphavirus nucleocapsid (N) proteins (Figure S4A), we observed that the anti-Semliki Forest virus (SFV) N antibody cross-reacts with VEEV N (Figure 7I). Western blot analysis of infected cell lysates revealed that the average cellular concentration of VEEV N reaches approximately 16.8 μM by 16 hours post-infection (hpi) (Figure 7I). Due to the nuclear localization of GFP-VEEV N reporter, we segmented the cytoplasmic volume of U2OS cells, and estimated the cytoplasmic concentration of VEEV capsid to be ∼6.5 μM at 16 hpi (Figure 7J). Using the GFP-N reporter system, we determined that a cytoplasmic concentration of approximately 0.22 μM is sufficient to inhibit SG formation (Figure S5B), indicating that this threshold is attainable during the early stages of infection. Furthermore, recombinant VEEV N protein purified from *E. coli* potently suppressed G3BP1-

RNA LLPS (Figure 7K). We noted the nuclear localization of VEEV N when expressed individually in cells. This localization was confirmed using both Flag- and HA-tagged constructs, as well as immunofluorescence assays for the tag-free construct (Figure S5C). However, during viral infection, immunofluorescence staining with the SFV N antibody revealed predominantly cytoplasmic localization of VEEV N, exhibiting both diffuse and punctate patterns (Figure S5D). This discrepancy may arise from the co-translational fusion of N with other viral structural proteins or from interactions with viral components that retain N in the cytoplasm. Notably, in cells expressing HA-tagged N, subsequent VEEV infection led to cytoplasmic relocalization of N (Figure S5E), indicating that viral infection actively modulates N’s subcellular distribution.

Collectively, these results reveal a conserved and functionally unified mechanism by which alphavirus capsid proteins antagonize stress granule formation, underscoring a fundamental strategy employed across the *Alphavirus* genus.

### 2.8. The SG modulation mechanism mediated by alphaviral capsid is distinct from that of SARS-CoV-2 capsid

In recent years, numerous studies have investigated the SARS-CoV-2 nucleocapsid N protein, which modulates SG formation and exhibits dose-dependent inhibition of SG assembly in human cells^52–54^. We could recapitulate the dose-dependent SG inhibition by SARS-CoV-2 N in cells (Figure 8A,B). The interaction between the IDR region of the SARS-CoV-2 N, centered on F17 residue, and G3BP1 is essential for SG inhibition. Although SARS-CoV-2 N also contains a positively charged RNA-binding motif (Figure 8A), mutation of F17 to alanine, disrupting G3BP1 interaction^55^, dramatically reduced its ability to inhibit SG assembly, indicating that the positive charges within SARS-CoV-2 N are insufficient for this function (Figure 8B,C). Instead, a peptide fragment of SARS-CoV-2 N encompassing F17, not the RNA-binding domain, is sufficient to inhibit SG formation (Figure 8B,C).

**Figure 8.**
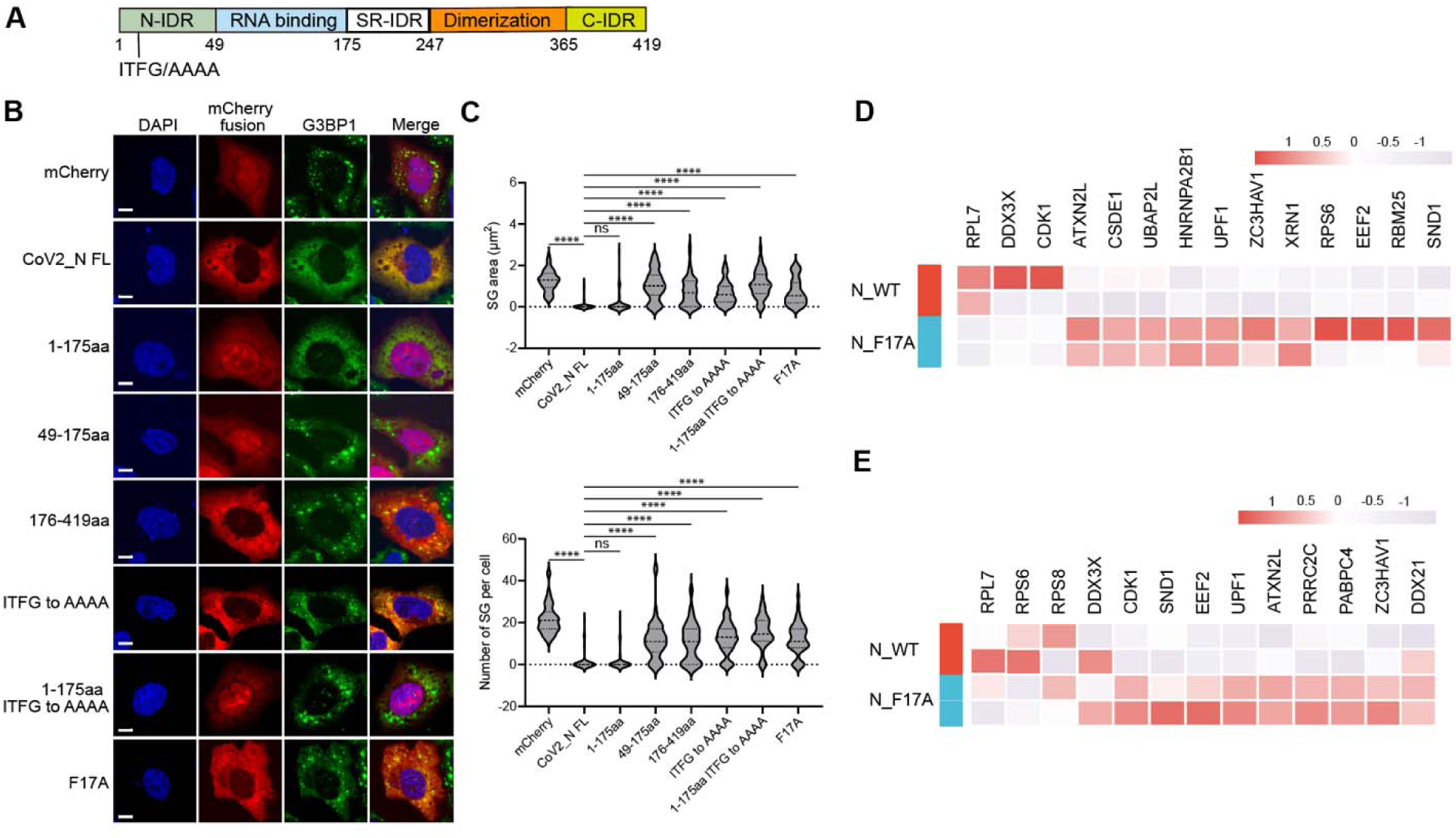
SARS-CoV-2_N inhibits SG via direct G3BP1 interaction. (A) Schematic illustration of SARS-CoV-2 nucleocapsid (N). A motif of ITFG and its alanine mutations was highlighted. (B) Immunofluorescent analysis of SA-induced SG formation in U2OS cells expressing the indicated mCherry-tagged mutants of SARS-CoV-2_N protein. Scale bar, 10 μm. (C) Quantification of SG area and number in (B). Data are plotted as minimum to maximum; lines indicate the first quartile (lower), median, and the third quartile (upper) with n≥50 for each sample (one-way ANOVA with Mixed-effects analysis). **p < 0.01, ***p < 0.001, ****p < 0.0001. Data are from one representative experiment for two independent experiments. (D, E) Heatmap of representative TurboID-G3BP1-enriched proteins in untreated (D) or SA-treated (E) U2OS cells co-transfected with SARS-CoV-2_N WT or F17A.

In contrast to the minimal impact of SFV capsid on the G3BP1 interactome, expression of SARS-CoV-2 N induces significant, F17-dependent remodeling of the G3BP1 interactome (Figure 8D,E). These findings, consistent with our own and previous studies, place SARS-CoV-2 N within the well-established mechanism of direct NTF2-like (NTF2L) domain binding and consequent G3BP1 interactome reorganization^55^. Therefore, our identification of alphaviral capsid-mediated SG modulation represents a mechanism distinct from that of SARS-CoV-2 N and its NTF2L-targeting pathway.

### 2.9. The SG-modulating feature of SFV N is not prevalent in viruses and the human proteome

All viruses possess a capsid for genome packaging. To assess the generality of viral capsid proteins in SG modulation, we synthesized a series of capsid proteins from diverse viral families, including vesicular stomatitis virus (VSV), dengue virus (DENV), Zika virus (ZIKV), and West Nile virus (WNV). We observed no significant inhibition of SG formation by VSV N either in cellular assays (Figure S6A,B) or *in vitro* (Figure S6C). Similarly, the N proteins of ZIKV, WNV, and DENV showed no substantial effect on SG assembly (Figure S6D,E). Notably, many of these capsid proteins localize to the nucleus. Given the prevalent nuclear-localizing features of New World alphaviruses, nuclear or nucleolar localization does not appear to be a determinant of SG inhibition. This observation argues against the involvement of a nucleolus stress pathway, potentially triggered by positively charged peptides, in SG suppression.

Using Foldseek^56^, a recently developed structural homology detection tool, we queried the helix-forming, charged region of the Semliki Forest virus (SFV) capsid as a template. This approach successfully retrieved homologous regions in other alphavirus capsids, including those of Chikungunya and Sindbis viruses (Figure S6F). However, most non-alphaviral capsids lack these conserved structural and electrostatic features, particularly the dense cluster of positively charged residues, indicating that the SG-inhibitory motif identified in alphaviruses is not widespread among other viral capsids.

Among the capsid proteins examined, HIV-1 nucleocapsid (NC) emerged as an exception, effectively suppressing SG formation (Figure S7A). NC is intrinsically disordered and contains a predicted short helical segment followed by two zinc finger RNA-binding domains. Expression of full-length HIV-1 Gag or the NC protein alone was sufficient to inhibit SG assembly. In contrast, mutations that disrupt the RNA-binding zinc fingers or neutralize adjacent lysine residues abolished this inhibitory activity (Figure S7B–D), suggesting that SG modulation by HIV-1 NC parallels that of alphaviral capsid proteins.

Given the enrichment of lysine residues in the functional motifs of these viral proteins, we investigated whether similar features exist in the human proteome. We analyzed the IDRome dataset curated by Tesie et al.^57^, derived from the UniProt human proteome (release 2021_04)^58^ and containing 28,058 IDRs. Based on pLDDT-based IDRome, we performed quantitative analysis of lysine abundance within each IDR. We identified 30 unique lysine-enriched IDRs, with lysine ratios ranging from 30% to 50% (Figure S8A). We then evaluated the effect on SG formation by expressing these lysine-rich IDR-containing proteins. To this end, we tested 22 proteins with available constructs, and none showed an effect on SG assembly or SG localization (Figure S8B).

Collectively, our sequence-based analysis and experimental validation indicate that the lysine-enriched, helix-prone motif capable of SG inhibition is rare in both viral capsids and the human proteome. This scarcity underscores the uniqueness of the identified motif and supports its potential utility in orthogonal design strategies for targeted manipulation of cellular processes.

### 2.10. SG modulators suppress stress granule formation induced by ALS-associated mutations

Stress granules (SGs) have been implicated in the pathogenesis of amyotrophic lateral sclerosis (ALS), in which aberrant phase transitions from liquid to solid states may contribute to protein aggregation, ultimately leading to neuronal cell death and disease progression^9,59,60^. Mutations in ALS-associated proteins such as FUS and hnRNPA1 are known to promote SG formation^59,61^. In this study, treatment with lysine-rich peptides significantly reduced FUS granule formation in FUS-expressing cells (Figure S9A,B). This suggests that the identified SG modulators may have therapeutic potential not only for neurodegenerative disorders but also for cancer and viral infections in which SG plays a pathogenic role.

## 3. Discussion

Here, we unexpectedly identified the inhibition of SG formation by the nucleocapsid (N) protein of Semliki Forest virus (SFV). This inhibitory function requires an oligomerization domain coupled with a lysine-rich segment (Figure 9, working model). Our findings indicate that this structural and functional feature is highly conserved across the alphavirus family but is absent in many other pandemic-prone viruses, suggesting a unique evolutionary adaptation specific to alphaviruses.

**Figure 9.**
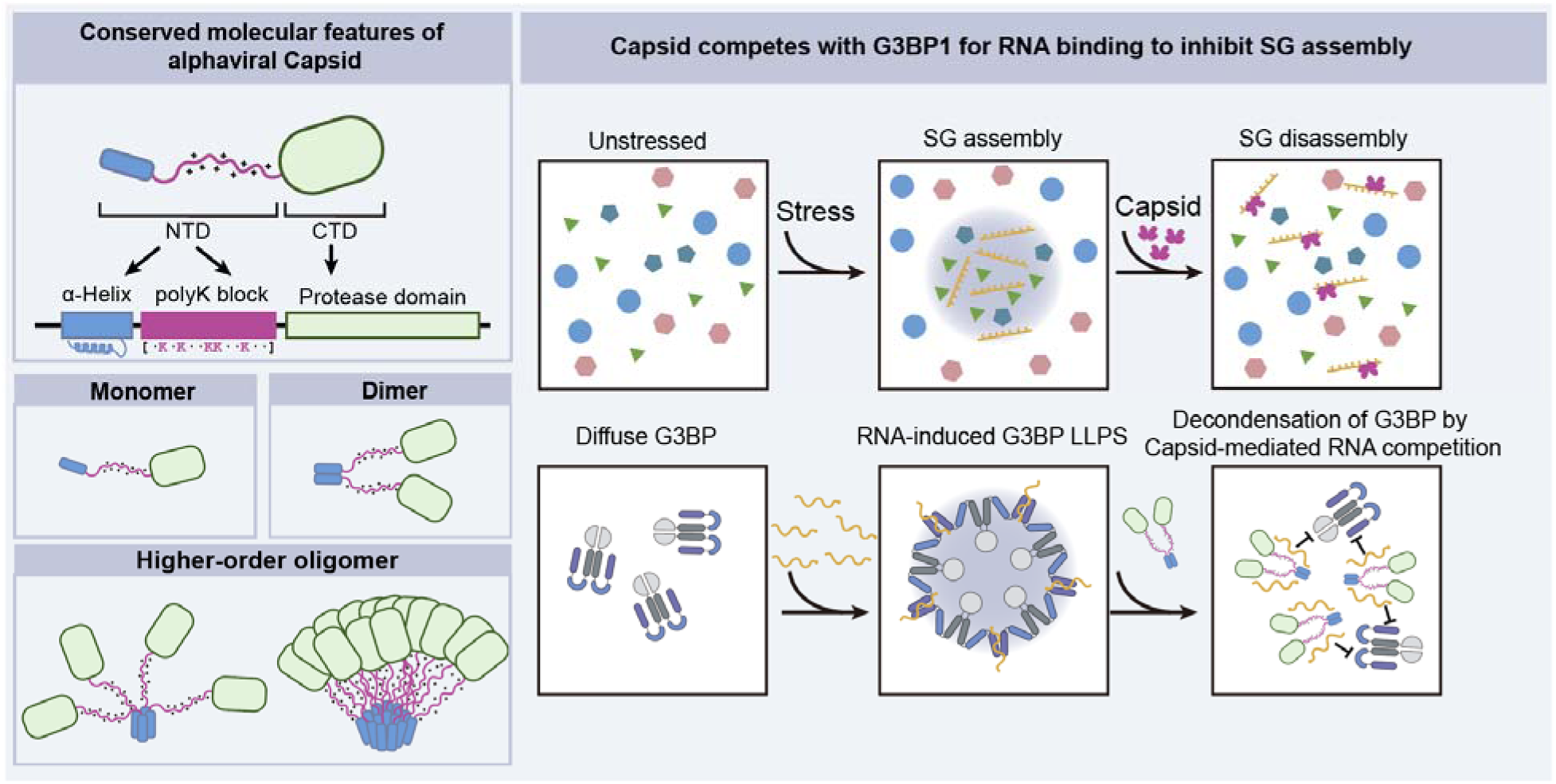
A working model. The alphavirus capsid harbors a conserved molecular feature, a helix-coupled lysine-rich region, that can efficiently inhibit SG formation by decoupling RNA from G3BP1.

Upon viral infection, host bulk translation is shut down, and stress granules form to limit viral replication^15^. However, the mechanisms by which viruses counteract this antiviral defense remain incompletely understood. Viral nucleocapsid proteins play critical roles in both genome packaging and modulation of host responses. For instance, the N protein of SARS-CoV-2 has been shown to impair double-stranded RNA (dsRNA)-induced PKR phosphorylation and to target G3BP1^62^, a key nucleating factor of SGs, thereby suppressing G3BP1-mediated SG assembly and enhancing viral replication^63,64^. Similarly, a recent study demonstrated that the N protein of Yellow Fever Virus directly interacts with the NTF2L domain of G3BP1 and delays the SG formation during virus infection^65^, mirroring the mechanism of SARS-CoV-2 N. Through comparative analysis, we demonstrated that the SG-inhibitory mechanism employed by SFV N is distinct from that of SARS-CoV-2 N.

While lysine-rich sequences are common across proteomes, we found that this specific combination, oligomerization capacity plus a lysine-rich motif, is not widespread, by screening the human proteome and diverse viral proteomes for similar features. This study uncovered lysine-rich sequences pairing with a short helix-forming segment within a compact peptide of fewer than 70 amino acids, which could exert potent effects on cellular RNA metabolism. RNA metabolism governs essential processes such as transcription, splicing, nuclear-cytoplasmic transport, translation, and mRNA decay. Stress granules, which assemble in response to infection, serve a broad antiviral role^66,67^. Our results demonstrate that the helix-coupled lysine-rich region efficiently suppresses SG formation. Disruption of the helical structure significantly impairs viral replication and correlates with increased SG formation in SFV-infected cells, supporting a functional link between helix integrity and SG inhibition. These findings further validate the long-predicted potential of coiled-coil helix formation in alphaviral capsid proteins^68–70^. Notably, an oligomerization-prone helix has also been recently identified in the IDR of the SARS-CoV-2 N protein^71^, although it doesn’t play a significant role in SG inhibition.

We propose that competitive RNA binding between SFV N and G3BP1 underlies the suppression of antiviral SG assembly. This mechanism aligns with emerging evidence that positively charged IDRs can act as RNA chaperones^72,73^. We hypothesize that the lysine-rich segment, when coupled with the oligomerization helix in alphaviruses, serves dual roles: facilitating efficient genome packaging and counteracting antiviral SG formation, particularly given its high expression during infection. Our data provide evidence for direct competition between SFV N and G3BP1 for RNA binding. Such a mechanism has recently been suggested for the SARS-CoV-2 N protein to prevent cGAS-RNA recognition^74^, as well as for herpesvirus ORF52 in restricting cGAS-DNA condensation^75^, and for MEX-5-mediated disassembly of the PGL-3 condensate^76,77^. Furthermore, the dependence of SG modulation on both the helix-forming region and the positively charged intrinsically disordered region (IDR) suggests a synergistic interplay among oligomerization, lysine valency, and RNA-binding capacity. The oligomerization potential of capsid proteins and their RNA-binding ability are likely mutually reinforcing and interdependent. Lysine residues may thus facilitate capsid protein oligomerization, while enhanced oligomerization could in turn increase RNA-binding affinity, and vice versa.

Dysregulation of RNA metabolism has been associated with various diseases, including tumorigenesis and neurodegeneration. Using the SG-modulating peptide identified here, we provide proof of concept for regulating aberrant SG dynamics induced by ALS-associated mutations. Unlike targeting protein-protein interactions directly on G3BP1 as reported in our studies (Yi Liu et al., unpublished), or small molecules that regulate cellular redox state to dissolve SG^78^, our approach highlights the therapeutic potential of modulating protein-RNA interactions.

Thus, our study identified a specific alphavirus-derived peptide capable of remodeling stress granules. Consistent with our discoveries, recent studies have demonstrated that various peptides can regulate biomolecular condensates, offering promising tools for bioengineering and therapeutic development. For instance, the Killswitch peptide, identified in a pathogenic frameshift variant of HMGB1, is hydrophobic and has limited homology to human peptides^40^. Killswitch has been used effectively to modulate nucleolar dynamics, oncoprotein condensates, and adenoviral nuclear condensates^40^. Small peptides composed of a specific combination of charged and aromatic residues have been shown to decelerate condensate ageing mediated by TDP-43 and FUS^79^. A peptide targeting the trimerization domain of oncofusion protein EML4-ALK disrupts its condensate formation and suppresses cancer cell proliferation^80^. Dipeptide repeats (DPRs) are translated via the repeat-associated non-AUG (RAN) mechanism from the G4C2 repeats within *C9orf72*^81,82^. Among them, positively charged peptides, including GR and PR repeats, modulate the dynamics of multiple biomolecular condensates and are major players in the pathogenesis of *C9orf72*-associated amyotrophic lateral sclerosis (c9-ALS)^81–83^. Together, these findings highlight a growing repertoire of biomolecular condensate modulators with significant potential for bioengineering applications and therapeutic interventions in disease.

The precise oligomeric state of the α-helix within the SFV capsid remains unresolved based on our CD analysis, crosslinking, and swap experiments. Future studies will determine whether this helix mediates dimerization, trimerization, or higher-order oligomerization, and whether the arrangement is parallel or antiparallel, parameters that may influence the spatial presentation of positively charged residues and their capacity to compete with G3BP1 for RNA binding. Elucidating the mechanistic details of this motif could enable the rational design of peptides with tunable SG-modulating activities, suitable for diverse physiological and pathological contexts. The conservation of this oligomerization helix across alphaviruses, combined with its essential role in viral replication, highlights its potential as a target for broad-spectrum, pan-alphavirus therapeutics.

## Figures and Figure legends

**Figure S1.**
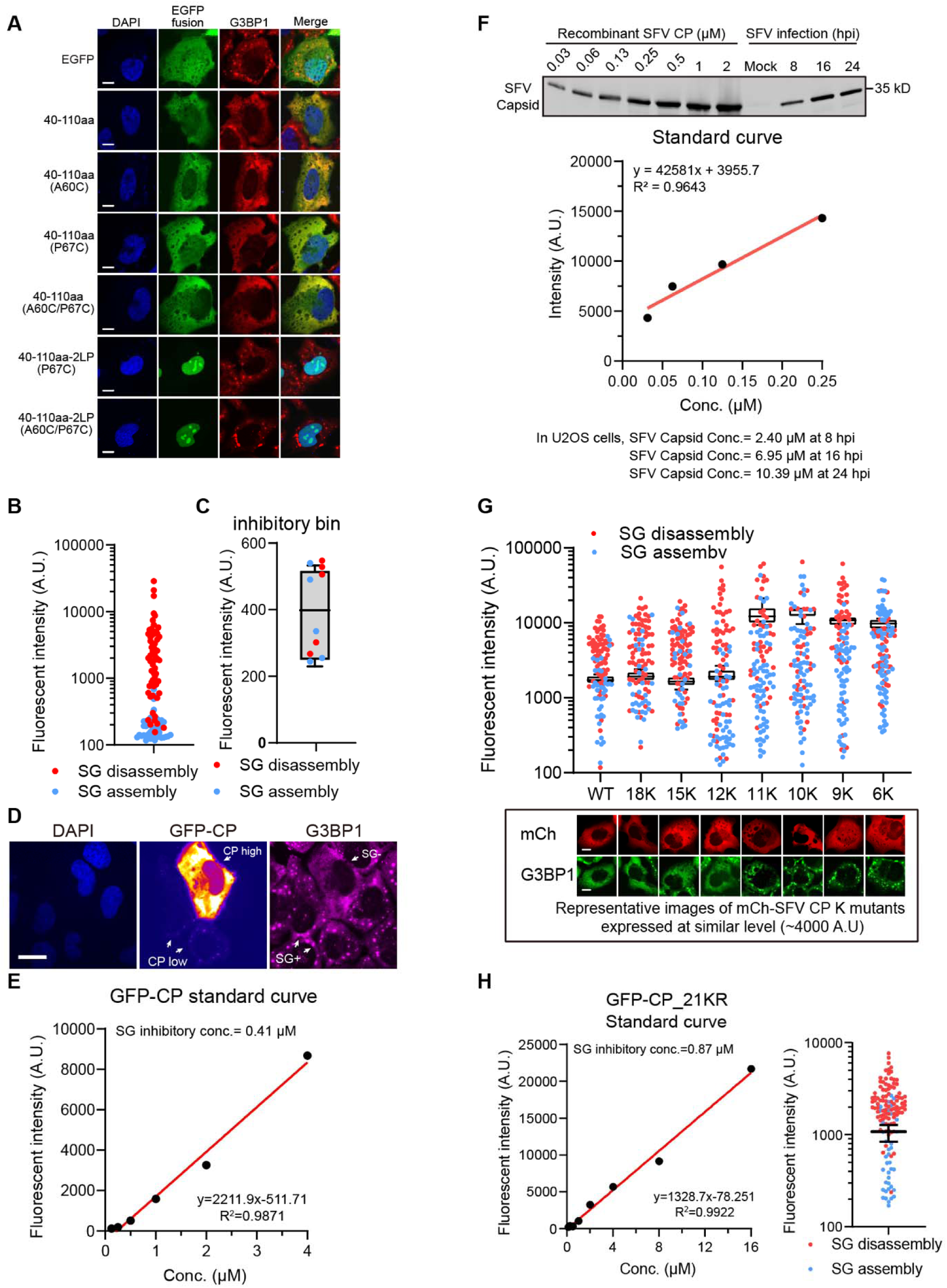
Determination of the threshold concentration of SFV Capsid in SG inhibition. (A) Immunofluorescent analysis of SA-induced SG formation in U2OS cells expressing the indicated EGFP-tagged cysteine mutants of SFV capsid-40-110aa. Scale bar, 10 μm. (B) Quantification of the relative fluorescent intensity of SFV capsid in SG inhibition. Data are plotted as a minimum to maximum. n=10 values in the determined thresholding bin. (C) The threshold concentration is shown. See Methods for defining the bin. (D) The representative image of SG disassembly in SFV capsid high-expressed cells, and colocalization of capsid with SG in low-expressed cells. Scale bar, 10 μm. (E) The standard curve of fluorescent intensity and transformation into the absolute SG inhibitory concentration of GFP-SFV capsid. The estimated SG-inhibiting threshold concentration of SFV Capsid is labeled above the curve. (F) Intracellular concentration estimation of total SFV capsid protein at 8, 16, and 24 hpi in SFV-infected U2OS cells (MOI=0.1). The estimated SFV capsid concentration is listed below the curve. (G) Quantification of threshold fluorescence of SFV Capsid-40-110aa lysine valency mutants in SG inhibition. Data are plotted as a minimum to maximum. n=10 values in the determined thresholding bin. Below are the representative images of the indicated mutants in 0.5 mM SA-treated U2OS cells. Scale bar, 10 μm. (H) The standard curve of fluorescent intensity and transformation into the absolute SG inhibitory concentration of GFP-SFV capsid-22-110aa-KR (left). The estimated SG-inhibiting threshold concentration of SFV Capsid is labeled above the curve. Quantification of the relative threshold fluorescence of SFV capsid-22-110aa-KR in SG inhibition (right). Data are plotted as a minimum to maximum. n=10 values in the determined thresholding bin.

**Figure S2.**
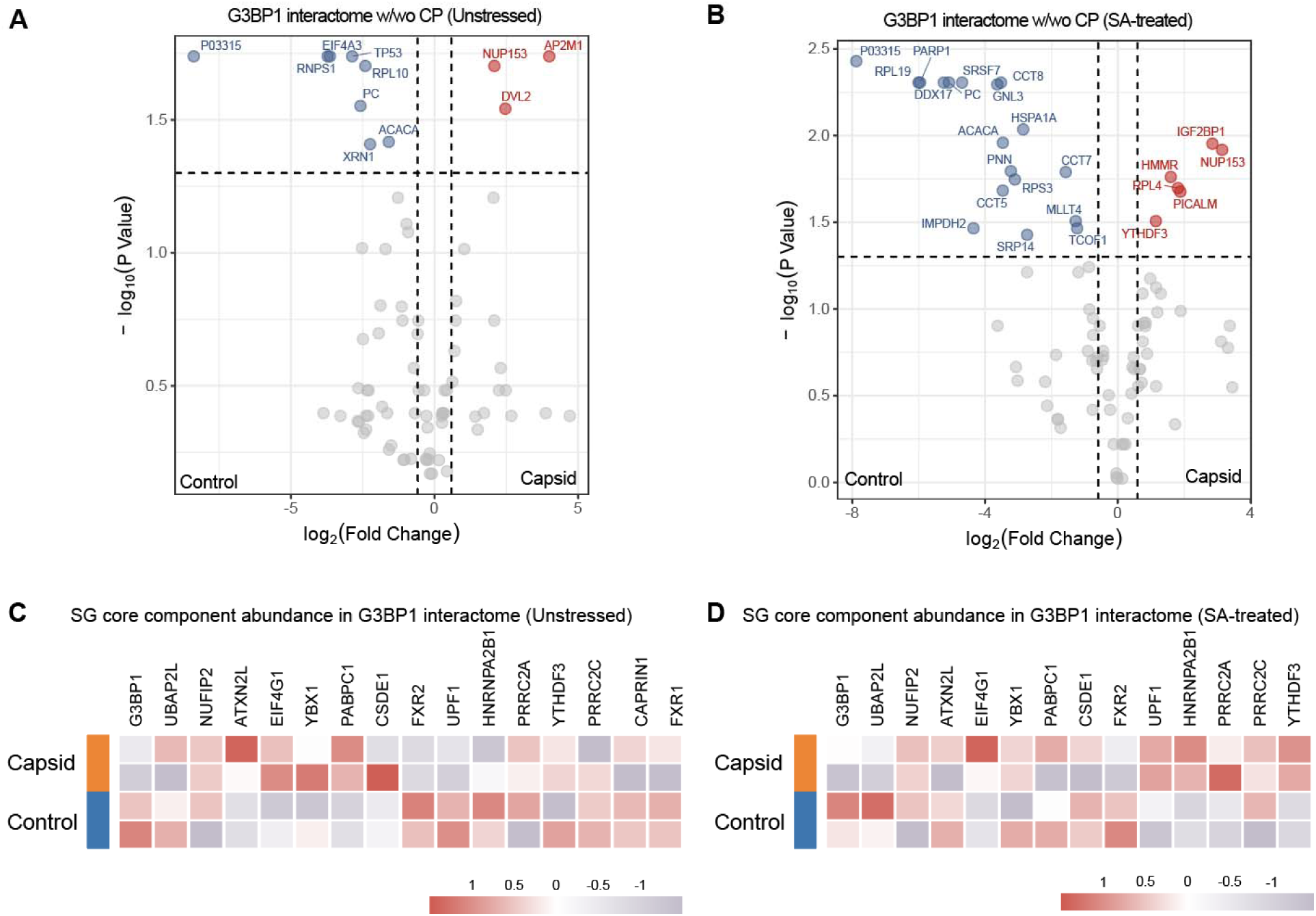
SFV capsid inhibited SG not by altering SG network. (A, B) Volcano plot of TurboID-G3BP1 proteome with or without SFV capsid in untreated (A) and SA-treated (B) HEK293T cells. (C, D) Heatmap of representative SG protein abundance with or without SFV capsid in untreated (C) and SA-treated (D) HEK293T cells.

**Figure S3.**
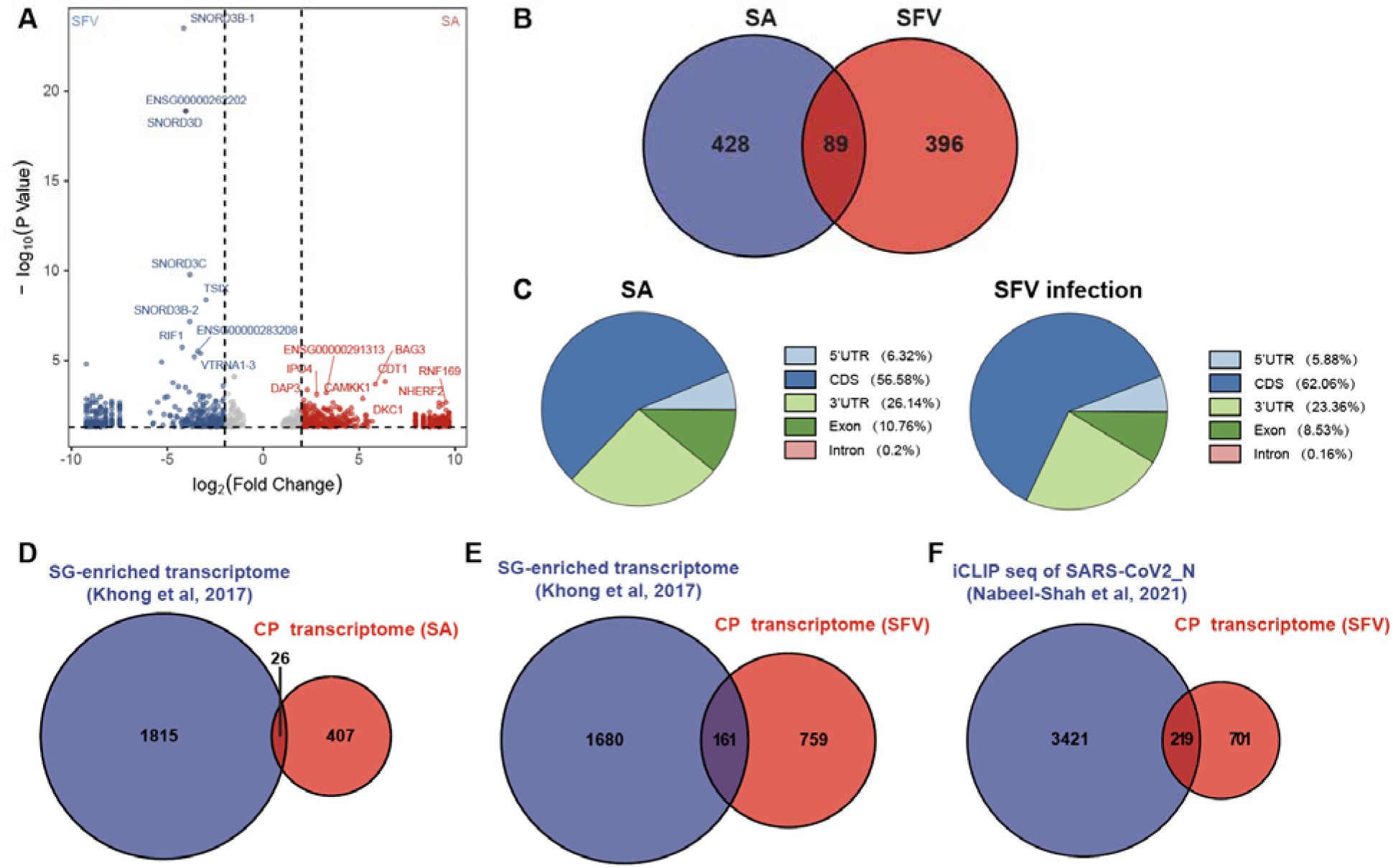
SFV capsid interacts with various host RBPs and mRNA. (A) Volcano plot of th binding transcripts of capsid in cells challenged with SFV and arsenite by capsid RIP-seq. (B) Venn diagram of SFV N-binding transcripts in cells challenged with SFV and arsenite. (C) Pie charts of capsid binding elements in the context of arsenite treatment (left) or SFV infection (right). (D, E) Venn plots of the SG-enriched transcriptome and the capsid binding transcriptome in SA-treated (D) or SFV-infected cells (E). (F) Venn plots of the SARS-CoV-2_N transcriptome and the SFV-infected capsid binding transcriptome.

**Figure S4.**
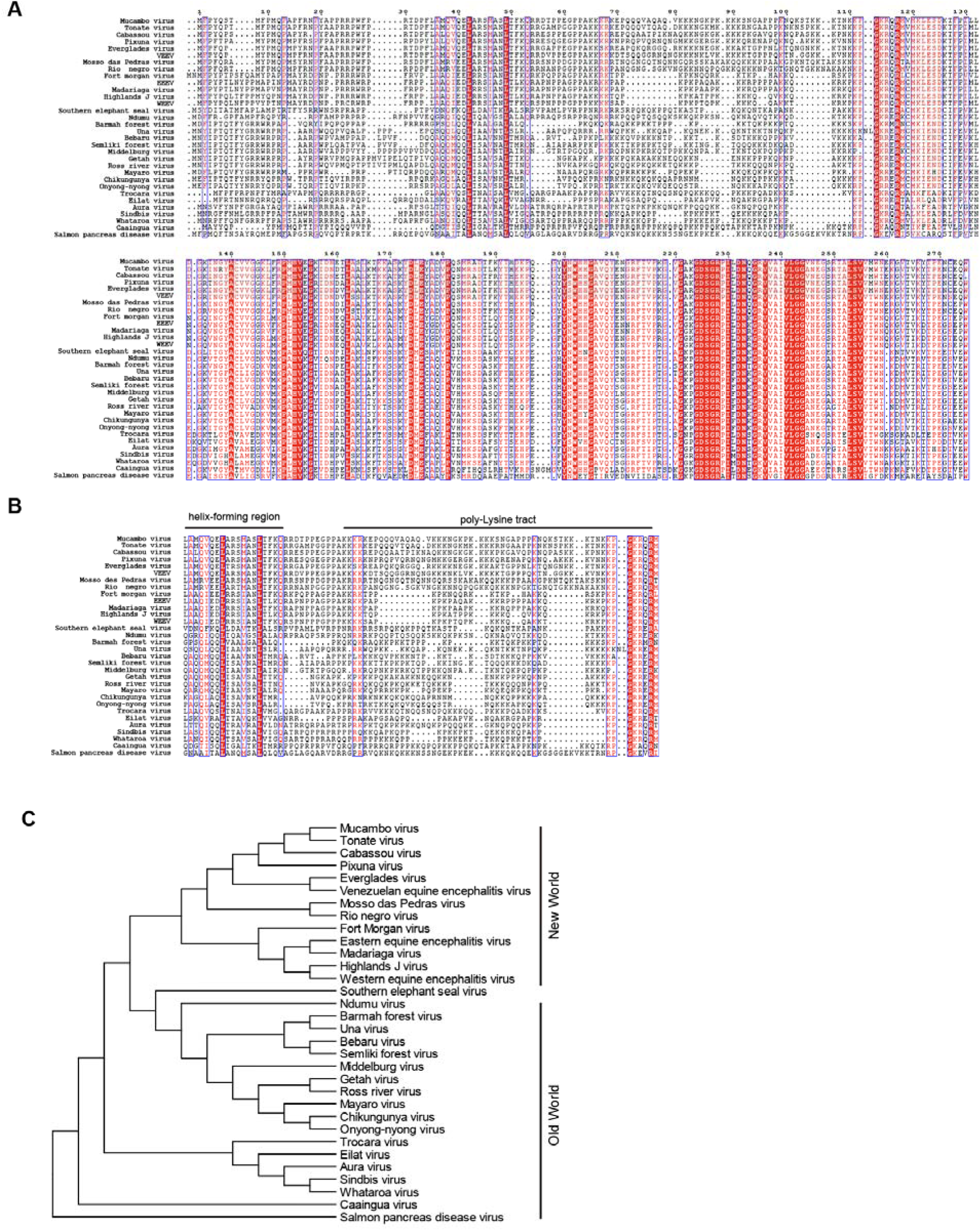
Amino acid sequence alignment of alphavirus capsids. (A) Alignment of capsid proteins of the alphavirus family members. (B) Alignment of helix-forming region and poly-lysine tract of alphaviral capsid proteins. (C) Phylogenetic tree of the alphavirus family based on the capsid protein sequences.

**Figure S5.**
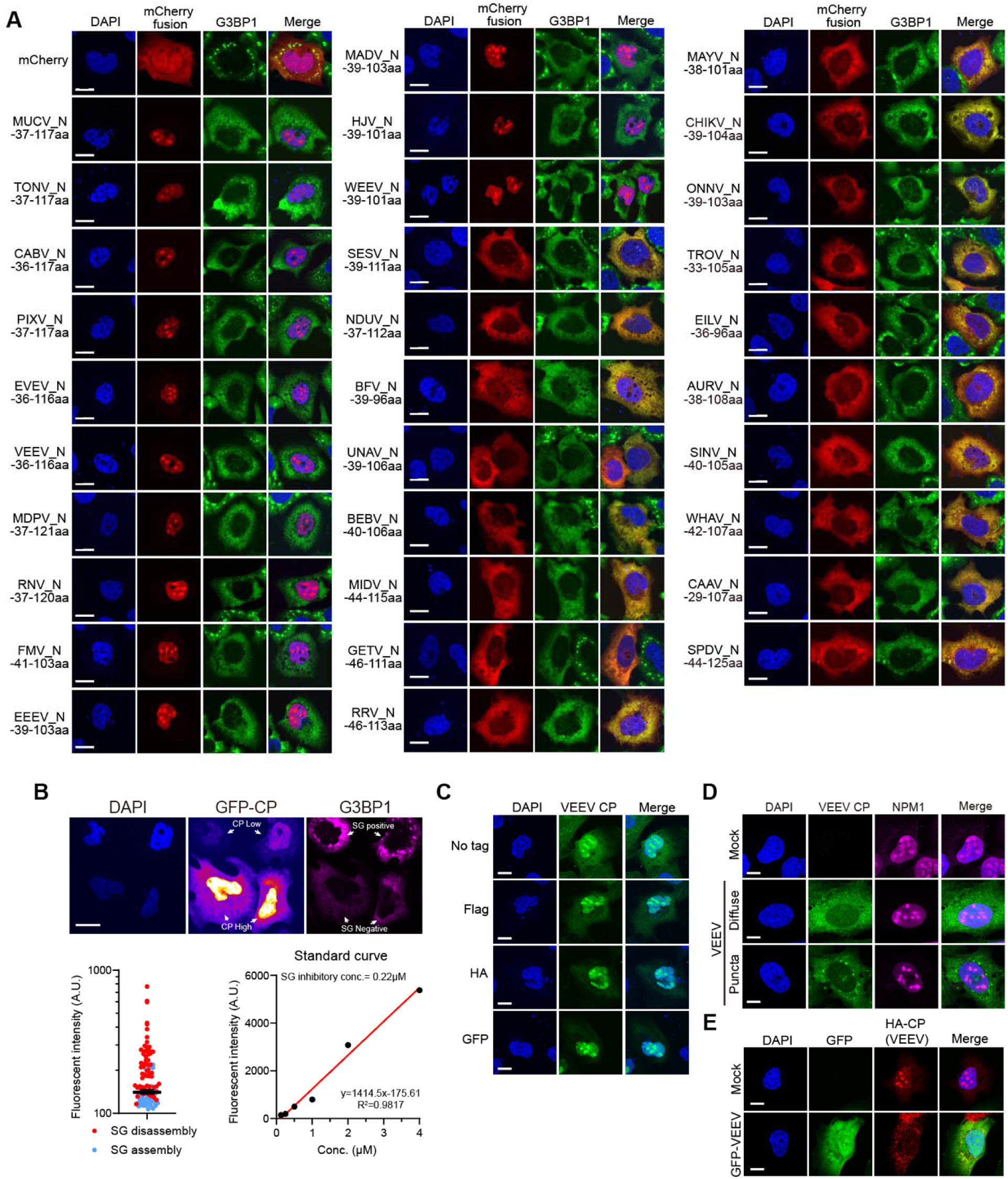
All alphavirus capsids showed SG inhibition. (A) The representative images of SA-induced SG formation in U2OS cells expressing the other 31 alphavirus capsid α-helix poly-lysine fragments. Scale bar, 10 μm. (B) The representative image of SG disassembly in VEEV capsid-expressing cells (top). Quantification of the relative threshold fluorescence of VEEV capsid in SG inhibition. Data are plotted as a minimum to maximum. n=10 values in the determined thresholding bin (bottom, left). The standard curve of fluorescent intensity and transformation into the absolute SG inhibitory concentration of GFP-VEEV capsid (bottom, right). The estimated SG-inhibiting threshold concentration of SFV Capsid is labeled above the curve. Scale bar, 10 μm. (C) Representative immunofluorescent images of VEEV capsids with different tags; the no-tagged VEEV capsid was stained with an SFV capsid antibody. Scale bar, 10 μm. (D) Confocal imaging of GFP-VEEV infected U2OS cells co-stained with NPM1 antibody. Two representative patterns were shown. Scale bar, 10 μm. (E) Confocal imaging of GFP-VEEV infected HA-VEEV capsid expressing U2OS cells. Scale bar, 10 μm.

**Figure S6.**
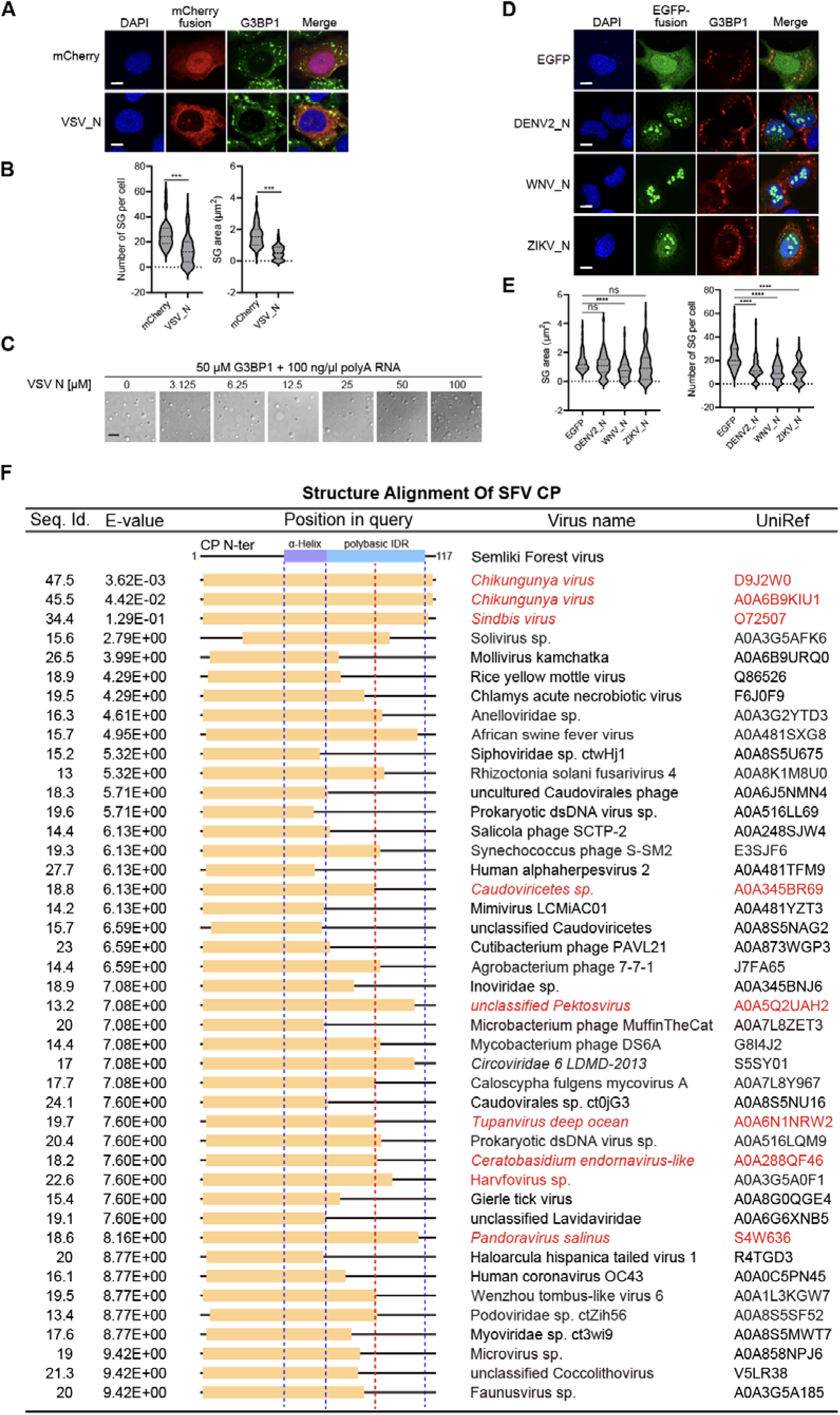
Many other viral nucleocapsids do not share the feature in SG inhibition. (A) Immunofluorescent analysis of SA-induced SG formation in U2OS cells expressing the mCherry-tagged VSV_N protein. Scale bar, 10 μm. (B) Quantification of SG area and number related to (A). Data are plotted as minimum to maximum; lines indicate the first quartile (lower), median, and the third quartile (upper) with n≥50 for each sample (one-way ANOVA with Mixed-effects analysis). ***p < 0.001. Data are from one representative experiment for at least two independent experiments. (C) The *in vitro* LLPS of 50 μM G3BP1 and 100 ng/μl polyA RNA in the presence of increasing concentrations of VSV_N. Scale bar, 10 μm. (D) Immunofluorescent analysis of SA-induced SG formation in U2OS cells expressing the EGFP-tagged DENV2, WNV, and ZIKV capsid proteins. Scale bar, 10 μm. (E) Quantification of SG area and number related to (D). Data are plotted as minimum to maximum; lines indicate the first quartile (lower), median, and the third quartile (upper) with n≥50 for each sample (one-way ANOVA with Mixed-effects analysis). ****p < 0.0001, ns is not significant. Data are from one representative experiment for at least two independent experiments. (F) The structure alignment of SFV capsid N-terminus with an online Foldseek tool (https://search.foldseek.com/search)^56^. The viral proteins sharing structural similarities with SFV are highlighted in red.

**Figure S7.**
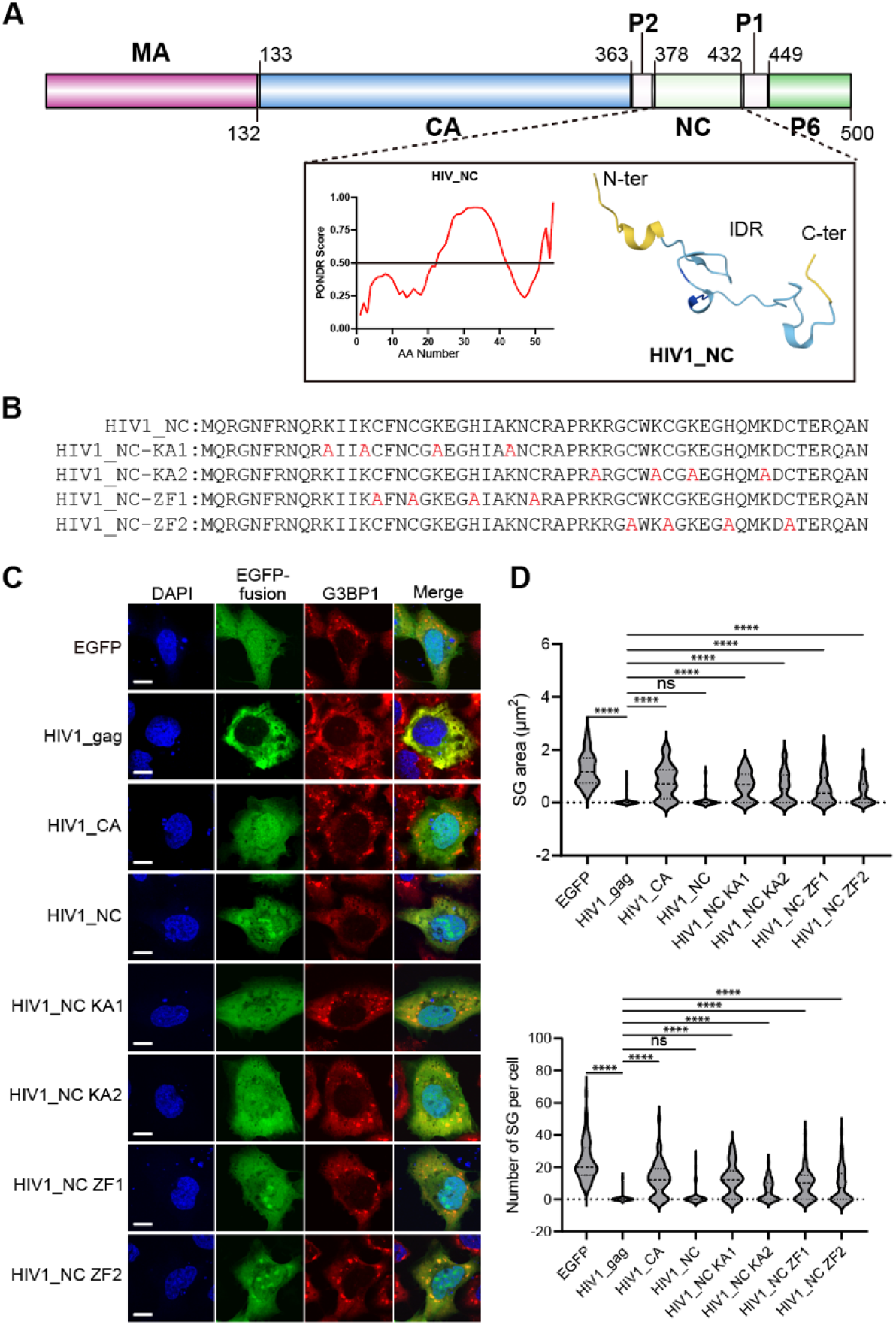
HIV-1 nucleocapsid inhibits SG formation. (A) Schematic illustration of HIV-1 gag protein domains, with PONDR analysis and Alphafold prediction of HIV-1 nucleocapsid. (B) Schematic illustration of the indicated HIV-1 nucleocapsid lysine to alanine mutants and ZF-disruptive mutants. (C) Immunofluorescent analysis of SA-induced SG formation in U2OS cell expressing the indicated EGFP-tagged HIV-1 gag, capsid, nucleocapsid protein WT, KA, and ZF mutants. Scale bar, 10 μm. (D) Quantification of SG area and number of SG related to (C). Data are plotted as minimum to maximum; lines indicate the first quartile (lower), median, and th third quartile (upper) with n≥50 for each sample (one-way ANOVA with Mixed-effects analysis). ****p < 0.0001, ns is not significant. Data are from one representative experiment for two independent experiments.

**Figure S8.**
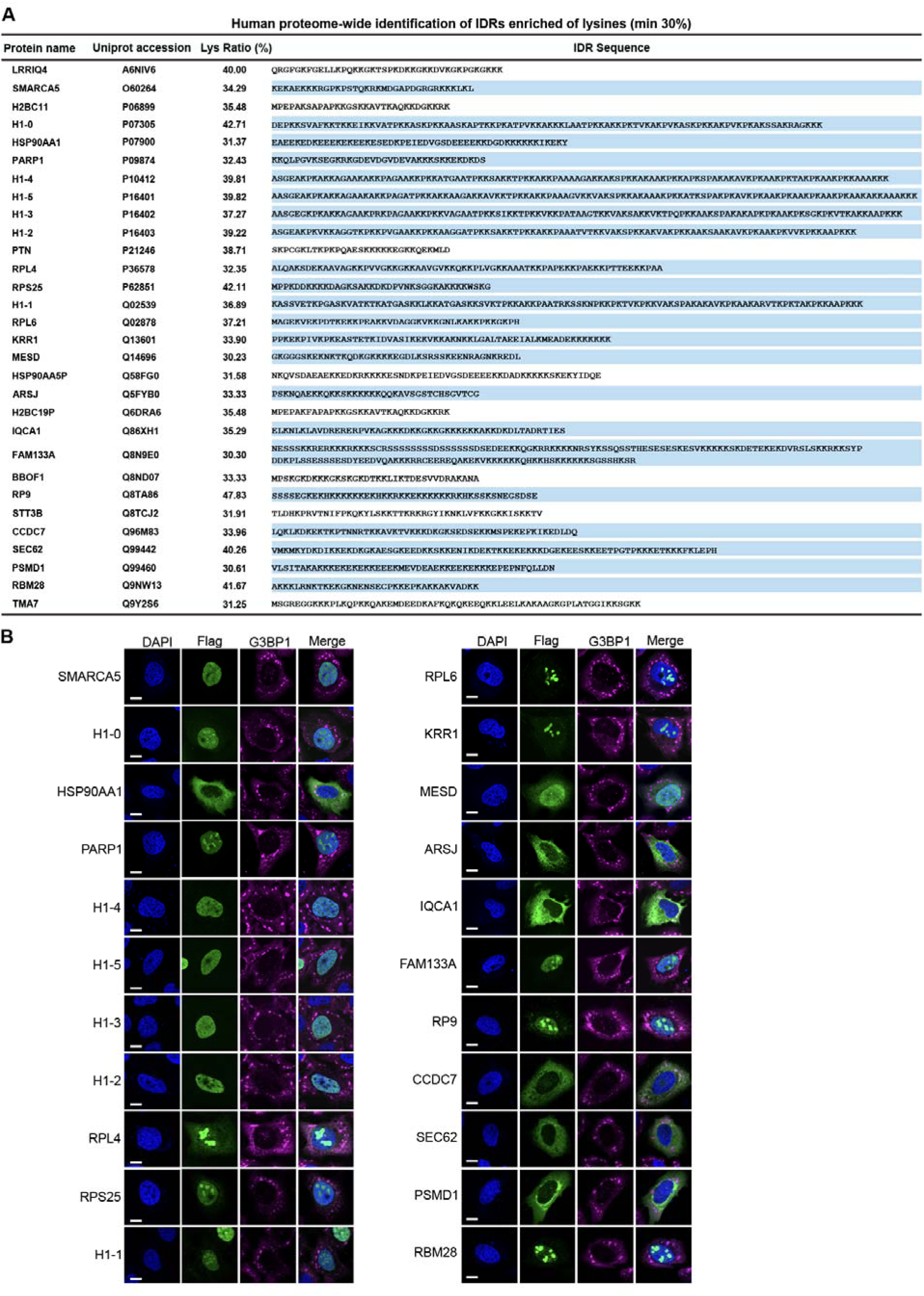
Proteome-wide search for capsid-like candidates. (A) A list showing 30 identified hits that contain an IDR with a lysine ratio over 30%. Highlighted genes are experimentally tested. (B) Representative confocal images of SA-induced SG formation in U2OS cells expressing the proteins with highly enriched lysine residues as listed in (A).

**Figure S9.**
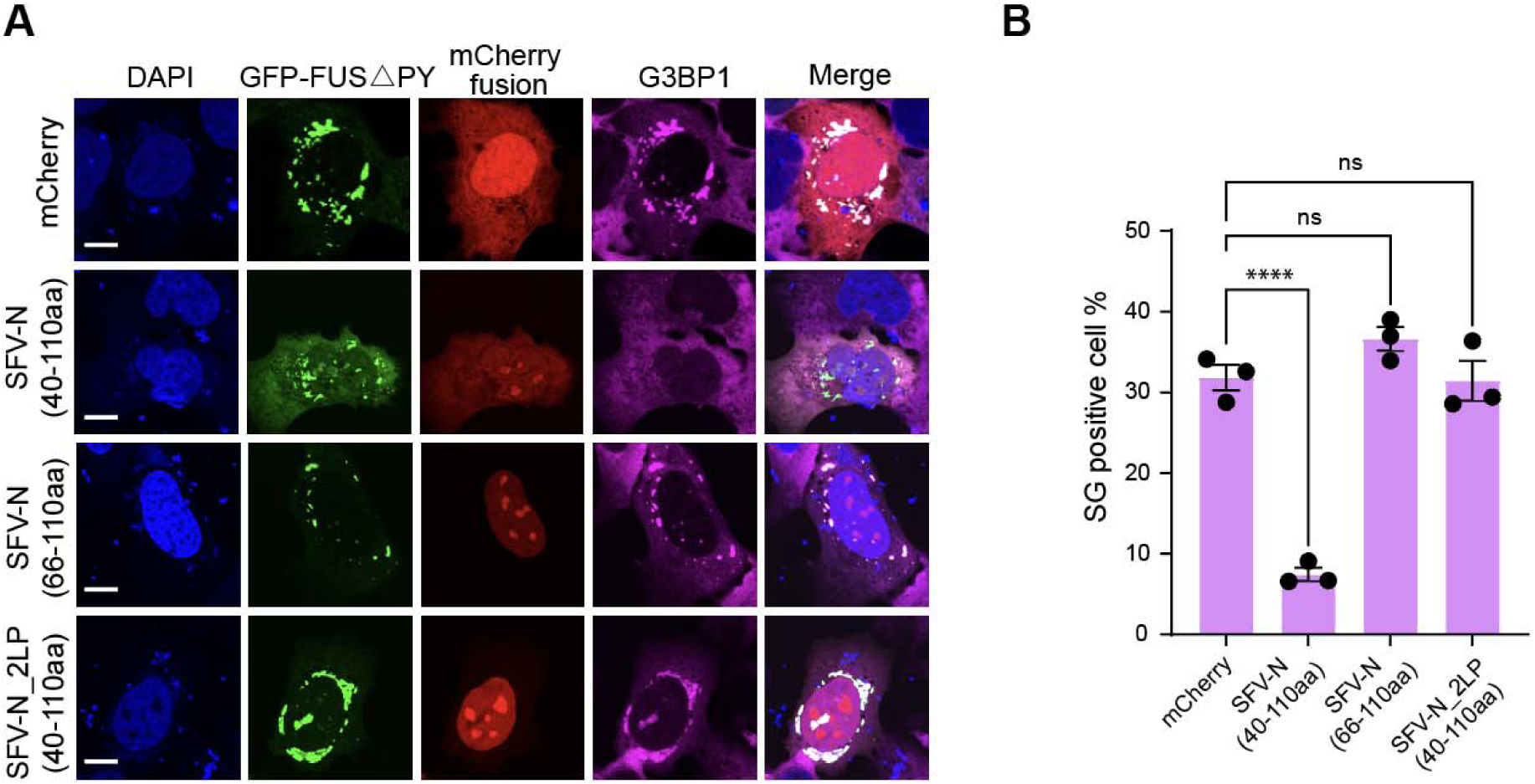
SFV capsid peptide inhibits stress granule formation induced by ALS-associated mutation. (A) Immunofluorescent analysis of FUS ΔPY-induced spontaneous SG formation in U2OS cells expressing the indicated mCherry-tagged SFV Capsid-40-110aa, 66-110aa, 40-110aa-2LP. (B) Quantification of cells with spontaneous SG in the indicated mCherry fusion-positive cells. Data are mean ± SEM with n=3 replicates. ****p < 0.0001, ns is not significant. Data are from one representative experiment for two independent experiments.

## 4. Methods

### Mammalian cell culture

The U2OS (RRID: CVCL_0042; #SCSP-5030), HEK293T (RRID: CVCL_0063; #SCSP-502), and BHK-21 (RRID: CVCL_1914; #GNHa10) cells were obtained from the Cell Bank, Chinese Academy of Sciences. All cells were verified as authentic by the provider via STR profiling. U2OS is used in cell biological studies due to its better cellular morphology for imaging. HEK293T cell is used in lentivirus packaging, and BHK-21 is used for SFV amplification. All cells were cultured in Dulbecco’s modified Eagle’s medium (DMEM) (Thermo Fisher, C11995500CP) supplemented with 10% Fetal Bovine Serum (FBS) (ExCellBio, FSP500) and 1% penicillin/streptomycin (Macklin, Q6532) and maintained at 37 ℃ in a humidified incubator with 5% CO_2_. Routine mycoplasma testing confirmed the absence of contamination in all cell lines.

### Bacterial cell culture

For plasmid amplification, *Escherichia coli* DH5α competent cells (TsingKe, TSC-C14) were used. Protein expression was carried out using BL21(DE3) chemically competent strains (TsingKe, DLC201). All bacterial cultures were grown in Lysogeny Broth (LB) medium (BD Biosciences, 214906) supplemented with the appropriate antibiotics at 37°C under constant agitation (200 rpm) in a shaking incubator.

### Plasmid construction

The sequences of SFV capsid/E1/E2/E3/E23/6K/nsP1/nsP2/nsP3/nsP4 were PCR amplified from pCMV-SFV4-INT construct and inserted into a modified pCDH-CMV-mCherry or EGFP vector for mammalian cell expression using the ClonExpress II One Step Cloning Kit (Vazyme, C112). The other alphavirus capsid used, SARS-CoV-2, VSV, DENV2, WNV, and ZIKV capsid, HIV1_ gag, SFV_N scrambled, 22-110aa-KR (21K within SFV capsid 66-110aa replaced with 21R) mutant, K^perfect^(1x), K(2x), K(3x), K(4x), K(5x), K(6x), and K(7x) (40-110aa with scrambled or shuffled sequence), 40-110aa-18K/15K/12K/9K/6K (66-110aa-with designated K replaced by S), were synthesized by General Biol with human codon-optimization and inserted into pCDH-CMV-mCherry or EGFP vector for mammalian expression. The coding sequences of G3BP1, SMARCA5, H1-0, HSP90AA1, PARP1, H1-4, H1-5, H1-3, H1-2, RPL4, RPS25, H1-1, RPL6, KRR1, MESD, ARSJ, IQCA1, FAM133A, RP9, CCDC7, SEC62, PSMD1, RBM28 and FUSΔPY (525-PY-526) were PCR-amplified either from the HEK293T cDNA or from a human whole genome ORF library (WZ Biosciences Inc, CH00001), and inserted into pCDH-CMV-mCherry or EGFP vector or pENTER-vector with a N-terminal flag-tag for mammalian cell expression. The HIV1_CA or NC coding sequences were amplified from HIV1_gag and cloned into the pCDH-CMV-EGFP vector. The SFV capsid mutants and truncations, including H145A, D167A, N-terminus, C-terminus, N-terminus-Δ Block1/ Δ Block 2/ Δ Block1+2, Block1, Block 2, Block1+2, 12-110aa, 22-110aa, 12-65aa, 22-65aa, 25-110aa, 28-110aa, 31-110aa, 34-110aa, 37-110aa, 40-110aa, 43-110aa, 46-110aa, 22-100aa, 22-90aa, 22-80aa, and the mutants and truncations of VEEV capsid, EEEV capsid, WEEV capsid, ONNV capsid, RRV capsid, SINV capsid, CHIKV capsid, SARS-CoV-2_N, HIV1_NC were constructed by one-step or two-steps reverse PCR from wild type sequence using Phanta Max Super-Fidelity DNA Polymerase (Vazyme, P505). The 2LP and 2LE mutations were introduced into the SFV capsid-encoding sequence at Leu45 and Leu52, respectively. The cysteine mutations were site-specifically introduced into the coding sequences of the SFV capsid fragment (40-110aa) and its 2LP variant, at the Ala60 and Pro67 residues, via one-step or two-step reverse PCR. The SFV_capsid 40-110aa-11K and 10K were subjected to mutation PCR from 12K and 9K, respectively. The HOTag3, HOTag6, Cry2, and Killswitch were inserted with a GSGS linker before the gene of interest. For the TetOn-EGFP/mCherry construct, the CMV promoter of the pCDH-CMV vector was substituted with a TetOn-inducible promoter, comprising 7xtetO elements and a minimal CMV promoter. The TurboID vector was constructed by inserting the TurboID-EGFP cassette downstream of the promoter in the modified pCDH-CMV vector. For *E. coli* expression, the SFV capsid full-length or truncations, VEEV capsid, GFP-SFV capsid full-length and 22-110KR mutant, GFP-VEEV capsid, and SARS-CoV-2_N were inserted between BamHI and EcoRI sites in a modified pMal-c5x-MBP-TEV-His backbone. And the VSV_N was cloned into a modified pET-32a-TrxA-His-TEV backbone. *E. coli* codon-optimized G3BP1 and G3BP1_EQ28 were reported previously^47^. All constructs were verified by bidirectional Sanger sequencing to ensure sequence accuracy.

### Protein purification

SFV capsid, N- or C-terminal of SFV capsid and 22-110-KR mutant, and VEEV capsid fused with N-terminal MBP-tag and C-terminal His-tag were expressed in BL21(DE3) cells. There is a TEV protease recognition sequence between the N-terminal MBP affinity tag and the C-terminal His-tagged target protein in the expression constructs. *E.coli* cultures were grown to mid-log phase (OD_600_≈0.8) and induced with 1 mM IPTG (JSENB, JS0154-100G) for recombinant protein expression at 16°C overnight. Cells were harvested by centrifugation and resuspended in ice-cold SEC buffer (50 mM Tris-HCl, pH 7.6, 400 mM NaCl) supplemented with protease inhibitors. Following resuspension, the cells were lysed by sonication on ice, and the crude lysate was clarified by centrifugation at 20,000g for 20 min at 4°C. The supernatant was subsequently loaded onto a pre-equilibrated maltose-binding protein (MBP) affinity column (5 mL bed volume of Dextrin Beads 6FF) (Smart-Lifesciences, SA026025), which had been pre-washed with lysis buffer at room temperature. Proteins were eluted with 10 mM maltose in lysis buffer. The eluted fusion proteins were subjected to tag removal by incubation with recombinant TEV protease in the presence of 0.1 mg/mL RNase A at 4°C overnight. The N- or C-terminal of SFV capsid proteins were precipitated without MBP fusion partner, we retained the MBP tag to enhance solubility and stability. The elution or cleavage reaction was subsequently purified by size-exclusion chromatography using a Superdex 200 Increase 16/200 column (Cytiva) pre-equilibrated with SEC buffer. Given the comparable molecular weights of the SFV capsid, VEEV capsid (∼35 kD), and GFP-SFV capsid-22-110aa-KR (∼40 kD) with MBP (∼40 kD), the elutions were further purified with Ni-NTA (Genscript, L00666). The MBP protein was eluted with 3 mM imidazole, and the target proteins were eluted with SEC buffer containing 250 mM imidazole. The purified proteins were replaced with SEC buffer and concentrated using 10 kDa MWCO Amicon Ultra centrifugal filter (Millipore, UFC801008), and stored at -80°C. G3BP1 and G3BP1_EQ28 proteins were purified with a fused GST-tag. *E.coli* cells were harvested, lysed, and centrifuged in SEC buffer. The proteins were purified with GST column and eluted with SEC containing 10 mM glutathione, removed tag with recombinant TEV protease, and treated with RNase A to reduce the nucleic acid contamination. The purified proteins were further applied to an equilibrated Superdex200 16/200 column in SEC buffer. SARS-CoV-2_N fused with MBP-tag and His-tag, and VSV_N fused with His-tag, were purified with Ni-NTA. The weakly-associated contaminant proteins were eluted with 30 mM imidazole in SEC buffer, and the objective proteins were eluted with SEC buffer containing 250 mM imidazole. The elutions were treated with RNase A, and the TEV protease was added into purified CoV2_N and VSV_N proteins. The elution or the post-cleavage sample was subsequently loaded on a Superdex200 16/200 column. SDS-PAGE was performed to analyze the fractions, and the target proteins were concentrated and stored -80°C.

### Immunofluorescence

U2OS cells were seeded onto coverslips in 24-well plates (Nest, 702001) and allowed to adhere overnight. The following day, cells were transfected with the respective constructs using polyethylenimine (PEI) (Polysciences, 24765-100) according to the manufacturer’s protocol. The culture medium was replaced with fresh medium 6 hours post-transfection, and the cells were then maintained at 37°C for 24 hours. For SG inhibition test, the transfected cells were treated with 0.5 mM Sodium Arsenite (Merck, 1.06277.1003) or 0.4M sorbitol for one hour at 37°C, or incubated at 43°C for 1 hour, then fixed with 4% paraformaldehyde (PFA) (Leagene, DF0135) for 10 minutes, permeabilized with 0.1% Triton X-100 in PBS for 10 minutes, and subsequently blocked with 1% bovine serum albumin (BSA) (Genview, FA016-25G) in PBST (PBS containing 0.02% Tween 20) for 1 hour at room temperature. The samples were incubated with primary antibodies diluted in blocking buffer at room temperature for 3 hours or 4°C overnight, followed by three 5-minute washes with PBST. Subsequently, the samples were incubated with appropriate fluorescent dye-conjugated secondary antibodies for 1 hour at room temperature, protected from light. For nuclear counterstaining and preservation, the slides were mounted using ProLong™ Gold Antifade reagent with DAPI (Invitrogen, P36931). Fluorescence images were acquired using an Olympus FV3000 laser scanning confocal microscope equipped with a 60× oil-immersion objective, with consistent acquisition parameters maintained across all experimental groups.

### Puromycin incorporation assay

For examining cellular bulk translation, cells transfected with mCherry-empty vector or mCherry-SFV capsid plasmid were treated with 10 μg/ml Puromycin (Biofroxx, 1299) for 15 min at 37 °C before being fixed with 4% PFA. Cells were further permeabilized, blocked, and incubated with an anti-Puromycin antibody (Sigma-Aldrich, MABE343) for immunofluorescence assay.

### 5’-ethynyluridine (EU) incorporation assay

To assess global transcriptional activity, cells were transfected with mCherry-empty vector or mCherry-SFV capsid plasmid. For the EU labeling, cell fixation, and Apollo staining, we followed the manufacturer’s instructions (Ribobio, R11081.1). Then the cell nucleus was stained with ProLong Gold Antifade reagent with DAPI (Invitrogen, P36931) for immunofluorescence assay.

### SG inhibition threshold concentration measurement in cells

U2OS cells were transiently transfected with GFP-SFV capsid/GFP-SFV capsid-22-110aa KR/GFP-VEEV capsid construct across a range of concentrations. Imaging was performed on a Nikon Eclipse Ti2 Spinning Disk Field Scanning confocal microscope equipped with a 60× Apochromat TIRF oil objective (NA 1.49) using identical capture settings. To build the GFP-SFV capsid concentration standard curve, recombinant GFP-SFV/GFP-SFV capsid-22-110aa KR/GFP-VEEV capsid protein was diluted two-fold, starting from 16 μM to 0.0156 μM. The fluorescent intensity of each capsid standard was recorded and plotted against the respective concentration to generate a calibration curve (Fig. S1E, S1H, and Fig. S5B). For each cell expressing the protein described above, a 2-μm diameter circular ROI was placed in the diffuse GFP region using ImageJ. Average intensity values from ≥100 cells were measured and converted to absolute concentrations via the standard curve. Cells were ranked by capsid protein concentration (high to low) and divided into 10 bins (10 cells/bin). The threshold concentration was defined as the mean concentration of the first bin containing >5 SG-negative cells (dispersed G3BP1 signal). Bin size-derived error was propagated for threshold estimates. The threshold concentration was further estimated based on the standard curve above. The use of a fluorescent protein standard curve in vitro for cellular estimation has been demonstrated previously^85^, due to the stability of GFP fluorescence under certain conditions^86,87^. Due to differences in the cellular environment and the *in vitro* GFP-fusion solution, as well as variations in pH and temperature, there may be a certain range of discrepancy between the estimated concentration and the actual threshold concentration. For relative SG inhibition threshold measurement of mCherry-SFV capsid K valency mutants, the indicated mCherry-tagged plasmids were transfected into U2OS cells for 24 hours. IF staining of G3BP1 was carried out, and the mCherry fluorescent intensities were quantified as above without further transforming to absolute concentrations.

### Quantification of SG area and number

Images were captured under consistent microscopy parameters (laser power, gain, resolution). The quantification was performed in ImageJ by first converting the image format to “RGB stack” and setting the scale. SGs in capsid-expressed cells were traced using the freehand tool. To enable quantification, images were subjected to thresholding with Max Entropy and adjusted to ensure each SG is segmented individually. The SG size and number in each cell were measured using the Analyze particle plugin. At least 50 capsid-positive cells were analyzed for each sample.

### Evaluation of SFV capsid-mediated inhibition of G3BP1_EQ28-assembled SG

To assess the ability of SFV capsid to suppress SG formation induced by the G3BP1_EQ28 mutant, G3BP1/2 dKO U2OS cells^47^ were co-transfected with EGFP or EGFP-SFV capsid, and mCherry-G3BP1 or mCherry-G3BP1_EQ28. Immunofluorescence assays were performed in three independent replicates. At least 50 EGFP/mCherry double-positive cells per replicate were imaged using an Olympus FV3000 laser scanning confocal microscope with a 60×oil-immersion objective under consistent acquisition parameters. SG-positive cells were defined as those harboring ≥3 G3BP1/G3BP1_EQ28 foci.

### Evaluation of spontaneous SG inhibition induced by ALS-associated mutations

GFP-FUS Δ PY and mCherry or mCherry-SFV capsid-40-110aa/ 40-110aa 2LP/ 66-110aa were co-transfected in U2OS cells. Immunofluorescence was performed to visualize G3BP1-marked spontaneous SGs. All experiments were conducted with three independent replicates. At least 50 GFP/mCherry double-positive cells per assay were captured using an Olympus FV3000 laser scanning confocal microscope equipped with a 60 × oil-immersion objective. All immunofluorescence imaging was performed under consistent acquisition parameters. Cells with spontaneous SG were defined as those with at least 3 G3BP1 foci.

### SFV capsid antibody generation

The rabbit polyclonal antibody against the SFV capsid was generated as follows: Purified His-SFV capsid protein served as the immunogen and was used to immunize two New Zealand White rabbits by HUAbio. Initially, the antisera containing SFV capsid were identified using Western Blotting. The SFV capsid antibody was then further purified from the antisera using Protein A/G. Finally, the functionality and specificity of the purified SFV capsid antibody were confirmed by Western Blotting.

### Lentiviral transduction and generation of stable cell lines

The GFP 1-10, GFP-TurboID-SFV capsid, TetOn-GFP-TurboID-control/G3BP1 constructs were co-transfected with packaging plasmids psPAX2 (Addgene, #12260) and pMD2.G (Addgene, #12259) into HEK293T cells using PEI. After 48 hours, lentivirus-containing supernatant was harvested and filtered through 0.45 μm PVDF membranes. For stable cell line generation, U2OS cells were transduced with the lentiviral medium for 1 hour in the presence of 10 μg/mL polybrene (Solarbio, H8761), followed by replacement with fresh complete medium. At 48 hours post-transduction, the top 10% GFP-expressing cells were isolated by fluorescence-activated cell sorting (FACS) using a Sony MA900 cell sorter. To induce protein expression, the TetOn-regulated cell lines were incubated with 1 μg/mL doxycycline (Solarbio, ID0390) for 24 hours before FACS sorting.

### Recombinant virus production and infection

The SFV genome cDNA construct (pCMV-SFV4-INT) used in this study was generously provided by Dr. Andres Merits (University of Tartu, Estonia). The construct contained a CMV promoter for mammalian expression, and a Rabbit β-globin intron was inserted in the capsid-encoding region for *E. coli* stability. To generate a recombinant SFV with GFP-tagged capsid, the GFP11-encoding sequences were inserted in-frame with the capsid coding region of pCMV-SFV4-INT and located between capsid residues Lys96 and Gln97^43^. For SFV-2LP/2LE, the 2LP and 2LE mutations were introduced into the capsid Leu45 and Leu52 residues in SFV genomic DNA construct via PCR. Infectious SFV particles (WT, GFP11-recombinant, or mutants) were produced by transfecting BHK-21 cells with the respective cDNA clones. Culture supernatants were harvested at 48 h post-transfection, cleared by centrifugation at 13,000g for 15 min at 4°C, and filtered through 0.45 μm PVDF membranes (Millipore) prior to aliquoting and stored at -80°C. For infections, cells were exposed to SFV (MOI=0.1-0.5) in serum-free DMEM at 37°C for 1 hour with intermittent gentle mixing, followed by culturing in complete medium until analysis. The GFP-tagged VEEV (TC83 strain) was kindly gifted by Dr. Wenchun Fan (Zhejiang University, China). For infections, both U2OS and HEK293T cell lines were inoculated at a multiplicity of infection (MOI) of 0.1.

### Plaque assay

To determine the viral titers, the BHK-21 cells were seeded in 12-well plates at 5×10^5^ cells/well and incubated overnight until 90–100% confluent. The viral stocks were 10-fold serially diluted with serum-free DMEM medium. Medium from 12-well plates was removed, and cells were washed once with PBS. 200 µL of each viral stock dilution per well was added in triplicate. The cells were incubated for one hour with gentle rocking every 15 min for adsorption. The medium was removed and washed once with PBS. 2 mL/well of overlay medium (1.5% agarose in MEM with 2% FBS) was added to immobilize virus and prevent secondary plaques. After additional 4 days’ incubation, the cells were fixed with 10% formaldehyde solution for 1 hour. After removing the fixative, the plates were rinsed with tap water and stained with 0.1% crystal violet (1 mL/well, 10 min). The plates were further gently rinsed with tap water to remove excess stain, air-dried, and counted for clear plaques under a bright light microscope (select wells with 20 – 100 plaques/well for accurate quantification). The viral titers were calculated using the formula: 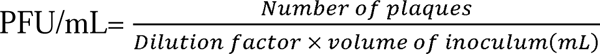

### Circular dichroism measurements

SFV capsid 35-64aa peptide (PDFQAQQMQQLISAVNALTMRQNAIAPARP) and 2LP peptide (PDFQAQQMQQPISAVNAPTMRQNAIAPARP) were synthesized by QYAOBIO, and measured by a Chirascan CD spectrometer (Applied Photophysics, UK) at room temperature. Samples were prepared in PBS to maintain proper folding conditions while minimizing signal interference from buffer components. Spectra were recorded at 190-260 nm with a step size of 1 nm and a cell path length of 1mm. The secondary structure quantification was performed using CDNN.

### Immunoprecipitation

For co-immunoprecipitation assays, cells were washed with PBS and lysed in IP buffer (20 mM Tris-HCl, pH 7.4, 150 mM NaCl, 1% Triton X-100, and 10% glycerol) supplemented with protease inhibitors for 20 min on ice, then centrifuged at 21,000g for 10 min at 4°C. 10% of the total lysate was saved as the input by denaturing in SDS loading buffer at 95°C for 10 minutes. For GFP/mCherry-tagged proteins, 1 μg/sample of homemade GST-GFP-nanobody or GST-mCherry-nanobody was conjugated to glutathione agarose beads (1 h, RT, rotation). The conjugated beads were incubated with lysates overnight at 4°C, then washed 3 times with IP buffer (10 min per wash, RT, rotation), and eluted by boiling in 2× SDS loading buffer. For proximity labeling assays, biotinylated proteins were lysed in RIPA buffer (20 mM Tris-HCl pH 7.4, 150 mM NaCl, 1% Triton X-100, 0.5% sodium deoxycholate, 0.1% SDS) with protease inhibitors. Biotinylated proteins were enriched with MyOne Streptavidin C1 Dynabeads (Invitrogen, 65002) at 4 °C overnight, washed once with 1 M NaCl and 3 times with RIPA buffer, and eluted by boiling in 2× SDS buffer containing 2 mM biotin.

### Western blotting

For routine assays, cells were lysed in RIPA buffer and 20 passes through a 1 ml syringe. Lysates or co-IP immunoprecipitants were separated on 4–20% gradient FuturePAGE gels (ACE Biotechnology, ET15420Gel) and transferred to nitrocellulose membranes (Pall). Membranes were blocked for 1 h at room temperature in TBST (Tris-buffered saline with 0.1% Tween-20) containing 5% (w/v) non-fat milk, then incubated with primary antibodies overnight at 4°C, and washed 3× with TBST (5 min per wash on an orbital shaker). Secondary antibody incubation was performed with IRDye 680RD/800CW (LI-COR; 1 h in the dark), followed by three additional TBST washes. Imaging was performed on a ChemiDoc MP system (Bio-Rad).

### Cysteine-mediated crosslinking

HEK293T cells were transfected with EGFP-tagged SFV capsid-40-110aa/40-110aa 2LP harboring cysteine substitutions and cultured for 24 hours, as described previously^39^. Subsequently, cells were rinsed with PBS and resuspended in ice-cold lysis buffer (50 mM Tris-HCl, pH 7.4, 150 mM NaCl, 1% Triton X-100, 1 mM EDTA) supplemented with protease inhibitors. Following 20 min of incubation on ice, cell lysates were centrifuged at 21,000 × g for 10 min at 4°C to pellet insoluble debris. The clarified supernatant was equally divided into two aliquots; one aliquot was supplemented with 5 mM DTT (Biofroxx, 1111GR025) to reduce disulfide bonds, while both aliquots were subjected to rotational incubation at 4°C overnight. The samples were denatured in 2× loading buffer (62.5 mM Tris-HCl, pH 6.8, 10% glycerol, 4% SDS, 0.05% Bromophenol Blue (Sigma Aldrich, BCCD6533)) at 75°C for 5 minutes. Denatured samples were then separated on 4-20% gradient FuturePAGE gels and transferred to nitrocellulose membranes for subsequent Western blot analysis.

### In vitro transcription

A DNA fragment containing a 5’-T7 promoter sequence (TAATACGACTCACTATAGGGAG) and the nsP3 coding sequence was amplified via PCR and purified using the FastPure Gel DNA Extraction Mini Kit (Vazyme, DC301). *In vitro* transcription was performed using 500 ng of the purified T7-nsP3 linear DNA fragment dissolved in RNase-free water with the T7 High Yield RNA Transcription Kit (Vazyme, TR101) and purified with the RNA Clean & Concentrator-5 kit (Zymo, R1015).

### Liquid-liquid phase separation assay in vitro

LLPS experiments were performed at room temperature unless otherwise indicated. Dilute protein stocks in 400 mM salt buffer to the desired concentration using 50 mM Tris-HCl (pH 7.4) to achieve a final salt concentration of 300 mM. Pre-mixed G3BP1 or G3BP1_EQ28 and polyA RNA (Sigma, 10108626001) or nsP3 RNA in a 0.2 mL PCR tube to induce phase separation. Add capsid/N proteins (SFV capsid, VEEV capsid, CoV2_N, truncated MBP-SFV capsids, or VSV_N) to the mixture, and the final salt concentration of buffer is 150 mM. Gently pipette the reaction mixture 3 times to ensure homogeneity. 2 μL of the reaction mixture was transferred into the observation chamber made of double-sided tape with a glass slide and a cover glass. All images were captured using a differential interference contrast (DIC) microscope with a 60×oil immersion objective within 5 minutes of reaction setup. For *in vitro* LLPS rescue assays, G3BP1 (25 μM) and polyA RNA (100 ng/μL) were pre-mixed with SFV capsid (12.5 μM) to suppress phase separation, followed by the addition of BSA, G3BP1, and polyA RNA at increasing concentrations.

### Estimation of intracellular protein concentration

For absolute intracellular SFV capsid and VEEV capsid concentration estimation, 5×10^5^ U2OS cells were infected with SFV or VEEV at a MOI of 0.1 (for Fig. 7I and S1F). Cells were harvested at the indicated time points post-infection. Infected cells were directly lysed in 50 μL of 2× SDS loading buffer to preserve protein integrity. Recombinant SFV capsid and VEEV capsid proteins were serially diluted by two-fold in 2× SDS loading buffer, generating concentrations ranging from 0.03 μM to 2 μM. 10 μL of each protein standard and cell lysates were separated on a FuturePAGE 4%–20% polyacrylamide gel. Proteins were transferred to a membrane and probed with a primary antibody specific to SFV capsid. Band intensities were quantified using ImageJ software. The average cell volume of U2OS was estimated to be 4×10^-12^ L^88^. The absolute intracellular protein concentration was calculated using the formula: 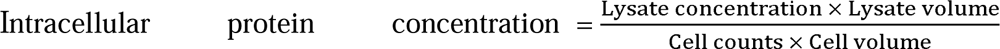

### TurboID proximity labeling

For the TurboID-mediated proximity labeling, the mCherry-control/SFV capsid/SARS-CoV-2_N WT/F17A were co-transfected with GFP-TurboID-G3BP1 in HEK293T cells for 24 hours before further treating with 500 μM SA for 1 hour. To initiate labeling, the culture medium was replaced with fresh medium containing 50 μM biotin (Sigma-Aldrich, B4501) for 10 minutes at 37°C. The labeling reaction was terminated by immediately placing the cells on ice and washing five times with ice-cold PBS.

### Fluorescence Polarization assay

G3BP1 and SFV capsid were serially diluted by two-fold (0-8000 nM) in 50 mM Tris-HCl (pH 7.4), 150 mM NaCl buffer, and 20 nM FAM-labeled RNA probe was added to each dilution. The sequences of RNA probes used are: FAM-AHNAK probe (5’-AUGCAAAGGGCUUGGACUUGGGUGGCAGAG-3’); FAM-GAPDH probe: 5’-UUUGUCAAGCUCAUUUCCUGGUAUGACAAC-3’; FAM-SFV gRNA probe1 (5’-GGUUGACGGGUCGGAUAGUUGCUU-3’); FAM-SFV gRNA probe2 (5’-GCGUCGUGUGAGCCCCCGAAAGACCACAUA-3’). The mixtures were incubated for 15 min at room temperature (protected from light) in black 384-well plates with clear bottoms (Beyotime, FCP987). All assays were carried out in triplicate. Fluorescence polarization (FP) was recorded using a BioTek Synergy Neo2 microplate reader, and polarization values were calculated using BioTek Gen5 software. The binding data were fitted to a one-phase association equation (nonlinear regression) in GraphPad Prism 10.

### RNA extraction and quantitative real-time PCR

Total RNA was extracted from cells using TRIzol reagent (Thermo Fisher, 15596018). Following RNA extraction, complementary DNA (cDNA) was synthesized using HiScript III RT SuperMix for qPCR (Vazyme, R312). Quantitative real-time PCR (qRT-PCR) was conducted on the Bio-Rad CFX Connect Real-Time PCR Detection System. The reaction was performed using Hieff qPCR SYBR Green Master Mix (No Rox) (Yeasen, 11201ES08) as the detection reagent.

### Identification of lysine-rich intrinsically disordered regions (IDRs)

IDRs were assigned briefly as follows: first, window-averaged AlphaFold2 pLDDT scores^89^ (window=20) were mapped to 20,215 human proteome sequences, and residues were categorized into three groups: folded (⟨pLDDT⟩ > 0.8), gap (0.7 ≤ ⟨pLDDT⟩ ≤ 0.8), and disordered regions (⟨pLDDT⟩ < 0.7).

To eliminate interference from fragmented and biologically irrelevant segments, folded or disordered regions shorter than 10 residues were reclassified as gap regions. Gap regions were subsequently reclassified based on their adjacent sequence context: those surrounded by disordered regions or located at the ends and surrounded by disordered regions were relabeled as disordered regions. All other gap regions were relabeled as folded regions. Finally, to ensure biological relevance, IDRs shorter than 30 residues or longer than 1,500 residues were excluded.

### RNA immunoprecipitation (RIP)

For determining RNA binding capacity of SFV capsid, HEK293T cells (2×10) were transfected with GFP-empty vector or GFP-SFV capsid for 24 h, followed by SFV infection for 12 h. Cells were washed twice with ice-cold PBS and a minimal amount of ice-cold PBS was added into cell dishes. Proteins and RNAs were cross-linked using a UV linker (V-leader, 150 mJ/cm²by energy mode). The cross-linked cells were lysed in RIP buffer (20 mM Tris-HCl pH 7.4, 500 mM NaCl, 10 mM EDTA, 0.5% Triton X-100, 0.1% NP-40, 1 mM DTT, 10% glycerol) supplemented with protease inhibitors (MCE, HY-K0010) and RNase inhibitor (Takara, 2313) in RNase-free tubes on ice for 30 min. Lysates were cleared by centrifugation at 13,000g for 10 min at 4°C. 10% of cleared lysate was mixed with 3 volumes of TRIzol reagent (Invitrogen, 15596026) and stored at −80°C as the input control. For immunoprecipitation, lysates were incubated with GFP-Trap Magnetic Agarose beads (ChromoTek, gtma) for 4 h at 4°C. Beads were washed 5 times with high-salt wash buffer (20 mM Tris-HCl pH 7.4, 500 mM NaCl, 10 mM EDTA, 1 mM DTT, 10% glycerol) containing protease inhibitor (5 min per wash, 4°C). Then the beads were resuspended in TBS containing RNase inhibitor, digested with 50 μg/mL Proteinase K (Promega, V3021) at 42°C for 15 min, The supernatant was collected and mixed with 3 volumes of TRIzol reagent to isolate the immunoprecipitated RNA and stored at −80°C.

### RNA-sequencing and data analysis

The input and immunoprecipitated RNAs from cells were stored in TRIzol reagent (Invitrogen) and extracted using Direct-zol RNA miniprep kits (ZYMO research) according to the manufacturer’s instructions. Genomic DNA was removed with DNase I and the RNA quality was verified with Nanodrop OneC spectrophotometer (Thermo Fisher Scientific) for purity and LabChip GX Touch system (Revvity) for integrity. The total RNA was applied to library preparation using KCTM Digital mRNA Library Prep Kit (Seqhealth Tech. Co., Ltd., Wuhan, China) according to the manufacturer’s protocols. The kit utilized a unique molecular identifier (UMI) of 12 random bases for labeling cDNA molecules, which can eliminate the duplication bias and errors produced in the PCR and sequencing steps. The quantification and sequencing of enriched libraries with 200-500 bp fragments were performed by the DNBSEQ-T7 platform (MGI) with PE150 sequencing mode. For data analysis, the RNA-sequencing reads were first filtered by fastp (v0.23.2), low-quality reads and adaptor sequences were removed. Clean reads were first clustered using UMI sequencing, followed by a new sub-clustering with > 95% sequence identity in the same cluster. For each sub-cluster, one consensus sequence was obtained by multiple sequence alignments. The processed sequences were mapped to reference genome GRCh38.110 using STAR software (version 2.7.6a). The gene expression levels were calculated with FPKM (Fragments Per Kilobase of transcript per Million mapped reads) by normalizing gene counts from featureCounts (version 1.5.1). The averaged FPKM of two biological replicates was used for the comparison. Differentially expressed genes were identified using edgeR package (version 3.40.2). A p-value cutoff <0.05 and IP over input fold-change cutoff >2 was applied.

### Mass spectrometry

For TurboID-mediated proximity labeling to identify the interactome of G3BP1, duplicate cell samples were subjected to streptavidin immunoprecipitation using magnetic beads. Proteins were separated on a FuturePAGE 4–20% polyacrylamide gel (160 V, 10 min). Gels with proteins were excised with a sterile single-edge blade, and processed for in-gel digestion. Gel slices were reduced with DTT, alkylated with iodoacetamide in 50 mM ammonium bicarbonate, and digested overnight with trypsin at 37°C. The peptides were separated using a Thermo EASY-nLC1200 nano-HPLC system coupled online to a Thermo Orbitrap Exploris 480 mass spectrometer. Separation was performed on a homemade fused silica capillary column (75 μm ID, 150 mm length; Upchurch, Oak Harbor, WA) packed with C-18 resin (300 A, 3 μm, Varian, Lexington, MA), with a 65-min gradient elution (flow rate: 0.3 μL/min). Mobile phases were 0.1% formic acid in water (A) and 0.1% formic acid in acetonitrile (B). The mass spectrometer operated in data-dependent acquisition (DDA) mode (Xcalibur 4.1 software), with a full MS scan (400–1800 m/z, 60,000 resolution) followed by 20 MS/MS scans (30% normalized collision energy). Data were processed using Thermo Xcalibur Qual Browser and Proteome Discoverer for database searching.

### Proteomic data analysis

For label-free quantification analysis of TurboID-G3BP1 proteome, raw data of each group were searched using MaxQuant software (v.2.4.9.0). The FASTA file containing the complete human proteome (downloaded from UniProt) as well as commonly observed contaminants was used as the reference for protein identification. The following search parameters were used: digestion was specified as trypsin with up to 2 missed cleavages. Fixed modification was carbamidomethyl[C] and variable modifications were oxidation of methionine and acetyl on protein N termini. Spectra were searched with a mass accuracy of 4.5 ppm for precursors and 20 ppm for fragment ions. The false discovery rate (FDR) was set to 0.01, both at protein and peptide levels, using a reverse database as a decoy. Label-free quantification (LFQ) parameters were set as default. For differential expression analysis, the processed files were analyzed using an online LFQ-Analyst tool (10.1021/acs.jproteome.9b00496) according to the detailed user manual. Briefly, the MaxQuant proteinGroups.txt files were uploaded, adjusted p-value cutoff was set to 0.05 and Log2 fold change cutoff was set to 0.59. The Benjamini Hochberg (BH) method was used for false discovery rate (FDR) correction. Quantified results were visualized using the Volcano tool in Hiplot Pro (https://hiplot.com.cn/). To achieve relative protein abundance, LFQ intensities were processed using Perseus. Briefly, the Maxquant-processed files were filtered based on at least 2 valid values for each row. The missing value imputation was carried out by replacing values from normal distribution. Z-score normalization was performed for each row, and the values were used to plot heatmap using Hiplot Pro (https://hiplot.com.cn/).

### Sequence alignment and phylogenetic tree of alphaviral capsid

All available alphavirus capsid protein amino acid sequences were retrieved from the NCBI database. Multiple sequence alignment was performed using the ClustalW algorithm implemented in MEGA 11 software. The aligned sequences were then used to construct a phylogenetic tree based on the maximum likelihood (ML) method with MEGA 11. The resulting alignment was exported in FASTA format and visualized using the web-based tool ESPript 3.065 to highlight conserved residues and structural features.

### Quantification and statistical analysis

Statistical analyses were conducted using GraphPad Prism 10. Tests included unpaired Student’s t test, one-way ANOVA, or two-way ANOVA, as specified in the respective figure legends. For qPCR data (bar plots), values are presented as mean ± SEM, with individual data points representing parallel replicates. The stress granule (SG) counts, area measurements, and inhibition thresholds (violin/box-and-whisker plots) display the first quartile (lower line), median (middle line), and third quartile (upper line), with whiskers extending from minimum to maximum values; each dot denotes a single cell. Significance thresholds: *\*p < 0.05*, *\*\*p < 0.01*, *\*\*\*p < 0.001*, *\*\*\*\*p < 0.0001*; ns (not significant).

## Acknowledgement

We thank the technical support from Yinping Gao, Fang Xiao, Fuli Zhao, and Yalin Wang from the imaging facility; Weiyuan Jin from the high-throughput screening platform; Jia Chen, Jin Hu, and Shan Feng from the proteomic center; and others at the Westlake Biomedical Research Core Facility. This work was supported by grants from the National Key Research and Development Project of China (2025YFC3409700 to P.Y.), Interdisciplinary Research Project of Hangzhou Normal University (2025JCXK02 to X.C.), the China Postdoctoral Science Foundation (2025M772812 to Z.Y.), the National Natural Science Foundation of China (32470733 and 32170696 to P.Y.), and the Center of Synthetic Biology and Integrated Bioengineering of Westlake University (WU2022A002).

## Author contribution

Y.Z. and Y.L. performed cellular assays. Y.Z., Y.L., and Z.Y. performed *in vitro* phase separation assay. H.L. and X.Z. performed the CD study and analyzed the data. X.C. and Y.Z. performed FP assay and analyzed the data. Z.Q. and Y.Z. generated the GFP10 cell and performed GFP11-SFV assay. Y.L. performed MS and RNA-seq studies and other related cellular and *in vitro* assays. Y.B. and T.L. analyzed the human IDRome and generated the list of K-rich proteins. P.Y., X.C., and X.Z. co-supervised the study. P.Y., Y.L., X.C., and X.Z. wrote the manuscript with input from all authors.

## Data Availability Statement

The mass spectrometry proteomics data have been deposited to the ProteomeXchange Consortium (https://proteomecentral.proteomexchange.org) via the iProX partner repository with the dataset identifier **PXD067786** (for the TurboID-G3BP1 interactome in the presence of SFV capsid) and **PXD067787** (for the TurboID-G3BP1 interactome in the presence of SARS-CoV-2_N_WT or F17A mutant). The raw sequence data regarding RNA binding profile of SFV capsid in untreated, SFV-infected, or Sodium Arsenite-treated HEK293T cells (**HRA012990)** reported in this paper have been deposited in the Genome Sequence Archive^90^ in National Genomics Data Center^91^, China National Center for Bioinformation/Beijing Institute of Genomics, Chinese Academy of Sciences that are publicly accessible at the following link: https://bigd.big.ac.cn/gsa-human/browse/HRA012990.

